# Parkin deficiency perturbs striatal circuit dynamics

**DOI:** 10.1101/636142

**Authors:** Magdalena K. Baaske, Edgar R. Kramer, Durga Praveen Meka, Gerhard Engler, Andreas K. Engel, Christian K.E. Moll

## Abstract

Loss-of-function mutations in the parkin-encoding PARK2 gene cause young-onset, autosomal recessive Parkinson’s disease (PD). Here, we investigated how parkin mutations affect cortico-basal ganglia circuit dynamics and cell-type-specific functional connectivity by recording simultaneously from motor cortex, striatum and globus pallidus (GP) in anesthetized parkin-mutant mice.

While ongoing activity of presumed striatal spiny projection neurons and their downstream counterparts in the GP was not different from controls, parkin deficiency had a differential impact on striatal interneurons: In parkin-mutant mice, tonically active neurons displayed elevated activity levels. Baseline firing of transgenic striatal fast spiking interneurons (FSI), on the contrary, was reduced and the correlational structure of the FSI microcircuitry was disrupted. The entire transgenic striatal microcircuit showed enhanced and phase-shifted phase coupling to slow (1-3Hz) cortical population oscillations. Unexpectedly, local field potentials recorded from striatum and GP of parkin-mutant mice robustly displayed amplified beta oscillations (∼22Hz), phase-coupled to cortex. Moreover, parkin deficiency selectively increased spike-field coupling of FSIs to beta oscillations.

Our findings suggest that loss of parkin function leads to amplifications of synchronized cortico-striatal oscillations and intrastriatal reconfiguration of interneuronal circuits. This presymptomatic disarrangement of dynamic functional connectivity may precede nigro-striatal neurodegeneration and predispose to imbalance of striatal outflow accompanying symptomatic PD.

## Introduction

Understanding early functional changes and adaptive mechanisms is a central aspect of modern research for Parkinson’s disease (PD) and key to the development of anti-parkinsonian treatment strategies at a preclinical level. During recent years, extensive studies in different genetic model systems have complemented conventional toxin-based dopamine depletion approaches and significantly advanced our pathophysiological understanding of familial PD from a molecular level to human clinical practice (Corti O et al. 2011; Klein C and A Westenberger 2012).

The prototypical example for genetically driven PD with autosomal-recessive inheritance is a loss-of-function mutation in the PARK2 gene encoding for parkin, a cytosolic ubiquitin E3 ligase (Kitada T et al. 1998; Lucking CB et al. 2000). While age of onset is earlier, compared to patients with idiopathic PD, the rate of disease progression in parkin-mutant patients is slowed (Khan NL et al. 2002; Pavese N et al. 2009) and dystonia is a frequent feature (Grunewald A et al. 2013). Parkin is required for efficient mitochondrial quality control and defective mitophagy is a candidate pathogenic mechanism for progressive nigro-striatal neurodegeneration in patients with parkin-associated PD (Scarffe LA et al. 2014). While much laboratory work is centered on the elucidation of patho-molecular mechanisms along the parkin pathway (Panicker N et al. 2017; Arkinson C and H Walden 2018), the consequences of parkin deficiency at a systems level are less clear. More recently, however, two separate lines of research have revealed important insights into the pathophysiology of parkin-associated PD.

Imaging studies in parkin mutation carriers provided evidence for a genetically driven presynaptic dopaminergic deficit early in the disease course (Hilker R et al. 2001) as well as subcortical and cortical changes in D2-receptor binding (Scherfler C et al. 2004). Moreover, heterozygote asymptomatic mutation carriers showed a mild pathology of the nigro-striatal pathway (Hilker R et al. 2001), an impaired sensorimotor information processing at a cortical level (Baumer T et al. 2007; Schneider SA et al. 2008) and enhanced cortico-striatal connectivity (Buhmann C et al. 2005). Moreover, similar to advanced idiopathic PD (Levy R et al. 2000; Brown P et al. 2001), network activity along the sensorimotor cortico-subthalamic axis of advanced parkin-associated PD is characterized by enhanced coupling in the beta frequency range (15-30 Hz) (Moll CK et al. 2015).

A second line of experimentation evolved in genetic small animal models, which do not typically display dopaminergic neurodegeneration and only partly reproduce clinical features of the disease (Dawson TM et al. 2010). Knockout models of the 3 proteins parkin, PINK1 and DJ-1 associated with autosomal recessive PD share a common pathway to regulate mitochondrial function (Dawson TM et al. 2010) and express only non-existent to mild behavioural changes (Goldberg MS et al. 2003; Itier JM et al. 2003; Madeo G et al. 2014). Parkin knockout mice at the age of 3-6 months are characterized by the absence of nigro-striatal cell loss and overt motor impairment (Itier JM et al. 2003; Meka DP et al. 2015). Dopaminergic cell loss as the pathological hallmark of PD is thought to be preceded by axonal loss of nigro-striatal projections and most likely occurs from a “dying back “ axonopathy (Raff MC et al. 2002; Dauer W and S Przedborski 2003; O’Malley KL 2010; Chu Y et al. 2012; Kordower JH et al. 2013), indicating that first neurodegenerative changes in PD take place at the nigro-striatal synapse before loss of dopaminergic cell bodies in the substantia nigra pars compacta. Parkin knockout mice hence open unique windows to study adaptive network changes at an early, premanifest disease stage without overt neurodegeneration.

In keeping with the interaction of parkin with proteins involved in synaptic vesicle metabolism (Sassone J et al. 2017), parkin knockout mice show mild alterations of nigro-striatal dopamine transmission—consisting of increased intraneuronal dopamine metabolism with increased oxidative stress in the striatum, reduced levels of dopamine and vesicular monoamine transporters (Itier JM et al. 2003) and decreased evoked dopamine release (Oyama G et al. 2010).

Furthermore, previous in vitro research has identified impaired dopamine metabolism and dysfunctional cortico-striatal synapses as pathophysiological hallmarks of parkin-associated (and other forms of monogenic) parkinsonism (Kitada T et al. 2009; Martella G et al. 2009; Madeo G et al. 2012).

How these two lines of research, however, converge at a mesoscopic network level is hitherto unclear—as the consequences of parkin deficiency have not been investigated at a systems electrophysiology level. We therefore set out to determine whether and how loss of parkin functions affects neuronal activity in cortex-basal ganglia circuits in vivo. To this end, we employed multi-electrode recordings in vivo to study adaptive electrophysiological circuit-level changes in parkin mutant mice.

## Materials and Methods

The present study was designed to characterize the impact of parkin mutations on neuronal activity along the cortico-striato-pallidal circuitry. To this end, we studied two groups of male parkin knockout mice (parkin^−/−^, n=8) and control littermates (parkin^+/+^, n=4) at a mean age of 160±15 days. In both experimental groups one animal died during anesthesia prior to electrophysiological recording sessions. Our mutant mice obtained an exon 3 deletion of the parkin gene (Itier JM et al. 2003) and loxP conditional alleles flanking exon 12 of the ret gene (Kramer ER et al. 2006; Meka DP et al. 2015). Thus, all animals carried floxed alleles of Ret (Retlx/lx) (Kramer ER et al. 2006; Kramer ER et al. 2007; Meka DP et al. 2015). Investigated parkin mutant mice did not show any significant dopaminergic neuronal loss in the midbrain as described previously (Itier JM et al. 2003; Meka DP et al. 2015). Two example Tyrosine hydroxylase (TH) stained sections with corresponding stereological TH-positive cell counts are shown in supplementary Figure 1. All experiments were approved by the Hamburg state authority for animal welfare (BUG-Hamburg) and were performed in accordance with the guidelines of the German Animal Protection Law.

### Animal preparation

Anesthesia was induced in a chamber filled with isoflurane. Following an initial injection of medetomidine (2 μg/10g body weight i.p.), mask ventilation with O_2_ and isoflurane (1-1.5 %) was initiated using a small animal ventilator (Inspira, Harvard Apparatus Inc., Holliston, MA) (Savola JM and R Virtanen 1991; Zuurbier CJ et al. 2002) and mice were then mounted in a custom-built small animal platform equipped with a stereotactic manipulator (Kopf Instruments, Tujunga, CA). Appropriate anesthesia plane was ascertained by the absence of tail and foot pinch reflexes, lack of movements and a continuous monitoring of vital parameters (body temperature, electrocardiogram, spontaneous breathing). Next, a craniotomy was performed on the left side of the skull (0.5-4mm lateral to midline, −2.5-+2.5mm anterior and posterior to bregma, respectively). Following the completion of surgery, target anesthesia level was achieved by lowering isoflurane concentration to 0.8±0.1%, complemented by low dose medetomidine (0.1μg/10g bodyweight). For termination of the experiment mice received a lethal overdose of ketamine prior to perfusion with 4% paraformaldehyde. Brains were removed from the skull and kept in a storage solution (30% sucrose for 2-3 days at 4°C).

### Histology and stereological cell counting

Tissue preparation and histological staining of dopaminergic (DA) neurons in the substantia nigra pars compacta (SNpc) and the ventral tegmental area (VTA) was performed according to the protocols of Meka (Meka DP et al. 2015) and Kramer et al. (Kramer ER et al. 2007). Midbrain cell count was performed for the right hemisphere by stereology (StereoInvestigator, MicroBrightField) and automatic fiber counting (Metamorph, Molecular Devices). To avoid counting biases, investigators were blinded to the genotype. The genotype was confirmed post-mortem with PCR. The left hemisphere was used to visualize recording sites. Brains were cut into 50 µm coronar sections with a freezing microtome (Leica Instruments GmbH) and fixed on slides. Sections were either stained with Nissl staining or Acetylcholinesterase (ACH) staining using standard protocols. Lesions and electrode trajectories were visualized using a microscope (Carl Zeiss Microimaging), photographed using AxioVision Software and merged using Fiji Software (Schindelin J et al. 2012). An example visualization of recording sites is shown in supplementary Figure 2.

### Data acquisition and experimental sessions

We recorded ongoing neuronal activity from 8 independently moveable microelectrodes (glass coated tungsten electrodes; tip size <5µm; tip impedances 670±240 kΩ; Alpha Omega, Nazareth, Israel) arranged in a 2×4 array (inter-electrode distance 0.8mm). During the course of the experiment, the microdrive was repositioned multiple times to cover a large extent of striatal tissue. Following the stereotactic positioning of the multi-electrode array, individual microelectrodes were lowered to the cortical surface in a stepwise fashion under microscopic guidance (EPS, Alpha Omega, Nazareth, Israel). Functional somatotopic mapping (somatosensory responses) at the cortical entry site allowed for a precise identification of the recording position and matching with atlas coordinates (Paxinos G and KBJ Franklin 2004). A single microelectrode was positioned as cortical reference within primary motor cortex. Depth of this cortical electrode was tailored towards robust unit activity and high signal noise ratio in superficial pyramidal cell layer II/III of M1, respectively. M1 was identified by both verification of atlas coordinates and the absence of brisk somatosensory responses to stroking the animals skin surface in different body parts. In contrast, the precise cortical entry position of neighbouring microelectrode tracks—lowered through somatosensory cortices—could reliably be verified by somatotopically distributed tactile responses.

The remaining 7 electrodes were lowered to the target structures in the caudate/putamen complex (CPu) and lateral globus pallidus (LGP), respectively. The final classification of the recording site was based on a combination of a) atlas coordinates, b) cortical mapping results, c) a stereotypical activity-depth profile of neuronal activities together with characteristic discharges of specific cell types (e.g., sparse spiking of presumed spiny projection neurons at a striatal level, tonic high frequency discharges of neurons within the LGP) along the trajectory, and d) post mortem verification of reference lesions at different depth levels. Recordings of uncertain location were excluded from further analysis. Supplementary Figure 2 illustrates the electrode configuration and verification of recording sites.

All signals were referenced against a silver wire positioned subcutaneously near the nose. Signals were pre-amplified, amplified (x 5000) and analogue-filtered (Multi Channel Processor; Alpha Omega). Spiking activity was extracted from the broadband signal (hardware filter settings 1Hz-12.5kHz) using a digital bandpass-filter between 160Hz and 6kHz and digitized with a sampling rate of 25kHz. Local field potentials (LFP) were obtained by applying a low-pass filter at 500Hz with a sampling rate of 1562.5Hz.

### Spike detection and sorting

All spike train analyses were carried out in Matlab (R2014b, MathWorks, Natick, MA). Spike detection and sorting was performed using the waveclus-toolbox (version 2.0) written by Quiroga et al. (Quiroga RQ et al. 2004). Threshold for spike detection was set at 4 times the standard deviation of the background noise of the bandpass-filtered signal. Action potential waveforms were aligned to the positive peak of the waveform. In this study, only well isolated single unit activity (SUA) was considered for further analysis. The selection criteria for SUA were twofold: signal to noise ratio had to be >2 and the summed standard deviation over the waveform’s main rise divided by the vertical length of the main rise had to be <3, according to the criteria described by Tankus and colleagues (Tankus A et al. 2009). Recordings with obvious drifts or artefacts were excluded from further analysis. Furthermore, units with less than 100 spikes were generally excluded from further analysis.

### Spike train analysis

As a first step of SUA analysis, we characterized the statistical spike train properties by calculating the interspike interval (ISI) distribution and the mean firing rate (defined as 1/mean ISI). To assess the regularity of neuronal discharges, we calculated both the global and local coefficient of variation (CV) of the interspike intervals. The local CV2 scales between 0 and 2, a value of 1 indicating Poisson-like properties and smaller values indicating more regular spiking (Holt GR et al. 1996). Given the inevitable activity fluctuations associated with prolonged recording durations, we selected the CV2 measure as a more accurate estimate of discharge regularity compared to CV. Neuronal group discharges (‘bursts’) were assessed with the Poisson surprise method described by Legendy et al. (Legendy CR and M Salcman 1985) and a surprise value of S=5.

### Classification of striatal neurons

The differentiation of striatal neuronal subtypes was a core aim of the present study. To this end, we used a multidimensional physiological parameter space to cluster our dataset, as described previously by Yamin et al. for the mouse striatum (Yamin HG et al. 2013). More specifically, we used both peak to peak width and half valley decay time (HDT) to separate presumed FSIs with short spike waveforms (likely corresponding to GABAergic interneurons) and presumed tonically active neurons (TANs, corresponding to striatal cholinergic interneurons) with long lasting afterhyperpolarizations (Bennett BD et al. 2000; Berke JD et al. 2004; Wilson CJ and JA Goldberg 2006; Sharott A et al. 2009; Lansink CS et al. 2010) from presumed spiny projection neurons of the striatum. These waveform measures were complemented by the proportion of ISIs longer than 2s, which has proven valuable in separating phasically active from tonically active neuronal striatal neuronal subpopulations (Schmitzer-Torbert NC and AD Redish 2008; Yamin HG et al. 2013). Unsupervised hierarchical clustering (McGarry LM et al. 2010) of the data was then performed using Ward’s algorithm (‘Exploratory Data Analysis’ toolbox (Martinez WL and AR Martinez 2005; Martinez WL et al. 2011)). Clustering results were visualized in a dendrogram and the number of groups was determined according to the results of the corresponding Mojena plot (Mojena R 1977). The quality of cluster separation was then assessed by calculation of the cophenetic correlation coefficient and a silhouette analysis (McGarry LM et al. 2010). Clustering performance was comparably good in both groups under study (mean silhouette values: parkin^−/−^ group, 0.72; control group, 0.73; average cophenetic correlation coefficient for the parkin^−/−^ group, 0.79; control group, 0.82). A cut-off value of 0.2 was used for the silhouette analysis in order to remove units with ambiguous cluster affiliation from further analysis (Kaufman L and PJ Rousseeuw 1990; Martinez WL and AR Martinez 2005). Therefore 38/290 (13%) of SUAs in the control group and 107/749 (14%) in the parkin^−/−^ group were excluded from further analysis.

### Correlation analysis

For statistical reasons, and in contrast to the descriptive spike train analyses above (firing rate, CV2, burst analysis), we only included single units with >300 spikes for all correlation analyses. Cross-correlations of simultaneously recorded SUA were computed in a time window of ±2 s with a bin size of 0.02 s (FieldTrip (Oostenveld R et al. 2011)). For statistical testing, ISIs of both spiketrains were shuffled 100 times using the global shuffling algorithm of Rivlin-Etzion (Rivlin-Etzion M et al. 2006). In order to correct for firing rate distortions, the mean null hypothesis was then subtracted from the true correlation. The null hypothesis was defined by the cross-correlation of the two shuffled spiketrains, and the mean null hypothesis was derived from the mean of 100 cross-correlations of the shuffled spiketrains. A t-score was calculated by dividing the corrected true correlation by the sum of the mean of all 100 standard deviations of the null hypotheses and the mean of the mean null hypothesis (Sharott A et al. 2009). Cross-correlograms were considered significantly modulated when >3 neighboring bins reached the significance criterion of ±2SD within a time window of ±0.25 s. The main peak of the cross-correlation was identified (defined as absolute value of positive or negative peak exceeding a minimum of >2SD within a central time window ±1.5s) and its time lag determined. In case both negative and positive peaks had similar heights, the peak closest to the central bin was chosen. The peak duration of cross-correlograms were determined by assessing zero-crossings next to the main peak. To further elaborate on the temporal relationships of correlated spiking, we calculated a symmetry index as in Adler et al. (Adler A et al. 2013). This symmetry index, defined as the difference of significant lags (>2SD) in the positive time range minus significant lags in the negative time range divided by their sum, was calculated for all t-scores in a central time window ±0.5s.

### LFP power analysis

Prior to spectral analysis, LFPs were down-sampled to 1000Hz and digitally band-pass filtered at multiples of 50 Hz to remove line noise with a second order notch filter. Moreover, we applied a linear phase low-pass filter at 200 Hz designed by the Parks-McClellan algorithm. All filtering procedures were applied by performing zero phase forward and reverse digital filtering. Power spectral estimates were calculated by using a multi-taper spectral approach (time-bandwidth parameter nw=3.5, 6 Slepian tapers, nfft=2000), resulting in a spectral resolution of 0.5 Hz. For group comparison, relative LFP power was retrieved by normalizing each spectrum by the total power between 1 and 80 Hz. In case more than 1 electrode was positioned within the recording structure (CPu, LGP) at a given time, power was averaged for all simultaneous recorded LFPs in the same anatomical structure before calculation of the group mean. Relative power was computed for five different frequency bands: delta (1.5-3.5Hz), theta (4-8.5Hz), alpha (9-19.5Hz), beta (20-30Hz) and gamma (30.5-80Hz; excluding the line noise region, 50±2Hz) and corrected for multiple comparisons using the false discovery rate. Delta- and beta frequency bands proved to be of specific interest in the context of this study. First, the high power values in the delta frequency band reflected the anesthesia-related low-frequency components. Secondly, mutant mice exhibited a distinct peak in the beta frequency range. Time-Frequency plots were constructed using the Chronux toolbox (Mitra PP and H Bokil 2008). For visualization purposes, we normalized the power spectra by dividing each spectrum by 1/f^1.5^as decribed by Feingold and colleagues (Feingold J et al. 2015).

### LFP coherence analysis

Coupling strength was determined by calculating the coherence *Cohij*(*f*) of two simultaneously recorded LFP signals *i* and *j* from different electrodes at a specific frequency *f*. This calculation is based on the cross-spectrum *Sij*(*f*), estimated using Welch’s averaged periodogram with a resolution of 0.5 Hz. The coherency *Cij*(*f*) can then be expressed as follows: 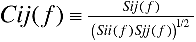 (Nolte G et al. 2004). To account for variance across recordings, coherency was transformed as suggested: 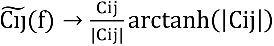 and the transformed coherence *Cohij* is then defined as the absolute value of the transformed coherency: 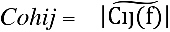 (Nolte G et al. 2004).

### Spike-LFP phase locking

Furthermore, we were interested in the precise temporal relationship between ongoing striatal and pallidal single cell discharges and population activities within and across these structures, respectively. Therefore, we calculated phase locking values of individual SUAs to simultaneously recorded LFPs from motor cortex, CPu and LGP. As a first step, LFPs were aligned to spike timestamps by downsampling to 1kHz. Similar to LFP power, phase locking of units to LFP was specifically assessed in the delta- and beta-frequency bands, respectively. To this end, LFPs were bandpass-filtered (Parks-McClellan filter) and Hilbert-transformed (Lachaux JP et al. 1999; Le Van Quyen M et al. 2001). As a measure of phase locking strength to LFP, mean phase and the vector length was derived from instantaneous spike field phases for each neuron (Sharott A et al. 2009; Sharott A et al. 2012). The significance of phase locking was determined by shuffling surrogate spike trains 1000 times (local shuffling of all ISIs in segments of 10s) and recalculation of the corresponding vector length. A neuron was considered significantly phase locked if the Raileigh statistic showed a non-uniform distribution with p<0.05 and more than 95% of the vectors obtained from the 1000 shuffled spike trains were smaller than the original one (Sharott A et al. 2009). LFPs were averaged at each depth in analogy to the spectral analysis described above. To discard possible influences of different discharge rates, we corrected for the number of spikes with a permutation procedure by 100 times randomly selecting 100 instantaneous spike-field phases for the calculation of the vector length.

### Experimental design and statistical analysis

A description of critical variables (number of recorded neurons, recordings and available data pairs) is provided in the results section. Due to non-uniformity of the data distribution, non-parametric statistical tests were generally applied. For pairwise comparisons, the Mann-Whitney U test (MWUt) was used. For comparison of categorical variables, Fisher’s exact test was used. For comparison of circular data (phase angles) we employed Williams-Watson-F test (WWFt) (Berens P 2009). P values were corrected with the false discovery rate (FDR) for multiple comparisons (Benjamini Y and Y Hochberg 1995). We corrected unless otherwise stated for each hypothesis for the number of compared neuron types, usually if available for 5 neuron types (4 striatal neuron types and 1 pallidal neuron type referred as LGP SUA). Significance level for all statistical tests was set at p<0.05 (individual p-values are given in the text). All given p-values in the results section are FDR-corrected, unless otherwise stated. All results are presented as mean±SD. Statistical analysis has been performed in MATLAB 2014b (Mathworks) and associated toolboxes, bar graphs and box plots have been created using Graphpad Prism. Whiskers of box plots show 5-95 percentile, outlier are for overview reasons not shown.

## Results

### Description of the dataset

We recorded ongoing neural activity simultaneously from motor cortex (henceforth referred to as Cx; 274 recordings), caudate putamen complex (CPu; 283 recordings) and lateral globus pallidus (LGP; 145 recordings) of isoflurane-anesthetized parkin mutant mice and a cohort of age-matched controls. Mean recording duration was similar for both groups (ctrl, 549±225s vs. parkin^−/−^, 564±237s; MWUt [n=228/474], *p*=0.69).

### Classification of different striatal neuronal subpopulations

A total of 1039 striatal single units, recorded along the whole anterior-posterior and medial-lateral axis of the CPu, passed our inclusion criteria (see materials and methods) and formed the starting point for further differentiation into different cell types. Similar to previous studies (Berke et al., 2004; Schmitzer-Torbert and Redish, 2008; Sharott et al., 2009; Yamin et al., 2013) we used a combination of waveform duration and discharge parameters to disentangle the physiological profile of different striatal neuronal subpopulations. As visualized in 3D scatterplots (based on peak-to-peak duration, half valley decay time and proportion of long ISIs of sorted single neurons), a similar pattern of three distinct clusters emerged both in the control group (Figure 1A; n=242) and the cohort of parkin mutant mice (Figure 1B; n=642). Established criteria (Wilson et al., 1990; Kawaguchi, 1993; Bennett and Wilson, 1999; Berke et al., 2004; Mallet et al., 2005; Schmitzer-Torbert and Redish, 2008; Sharott et al., 2009; Lansink et al., 2010; Adler et al., 2013; Prosperetti et al., 2013; Yamin et al., 2013; Thorn and Graybiel, 2014) allowed a robust allocation of these clusters to striatal neuronal subpopulations of spiny projection neurons (pSPN), fast spiking interneurons (pFSI) and tonically active neurons (pTAN). The prefix ‘p’ stands for *putative*, reflecting the inherent uncertainty of relating single cell recordings to a particular neuronal suptype in the absence of post-hoc verification (labelling). As expected, pSPNs outnumbered all other neuron types and accounted for ∼40% of all classified neurons in both experimental groups (ctrl, n=121; parkin^−/−^, n=324). pSPNs were characterized by a high proportion of long ISIs (indicating a phasic discharge mode), largely contributing to their low ongoing discharge rate, and an intermediate waveform duration (Figure 1F). pTANs, likely corresponding to the population of striatal cholinergic interneurons, made up ∼20% of the total sample (ctrl, n=63; parkin^−/−^, n=154). The distinguishing features of this cell class were the absence of long ISIs (>2s) together with long spike waveforms (Figure 1D). In contrast, the short and rather symmetric action potential waveforms were an outstanding feature of pFSIs, presumably corresponding to the parvalbumin-positive fast firing subpopulation of striatal GABAergic interneurons (Berke JD et al. 2004; Tepper JM and JP Bolam 2004; Sharott A et al. 2009; Lansink CS et al. 2010; Wiltschko AB et al. 2010). Notably, this short-waveform-based cluster scattered along the whole ‘long ISI proportion >2s’-axis in both groups under study, with a substantial fraction of cells with short waveforms exhibiting discharge attributes similar to pSPNs. In agreement with the classic distinction between phasically active neurons and those with much higher discharge rates (Schmitzer-Torbert and Redish, 2008) and to comply with a similar subdifferentiation recently suggested by Yamin et al. (Yamin et al., 2013), we further subdivided this cluster at a cut-off of 0.5 based on the bimodal distribution of the discharge parameter ‘proportion of long ISIs > 2s’. Thus, highly active cells with short spike waveforms (<50% of spikes spent in long ISIs) constitute the cluster of pFSIs (ctrl, n=56; parkin^−/−^, n=117; Figure 1C). A smaller fraction of cells with short spike waveforms firing sporadically (>50% of spikes spent in long ISIs) were designated as unidentified neurons (UIN; ctrl, n=12; parkin^−/−^, n=47; Figure 1E).

**Figure 1.**
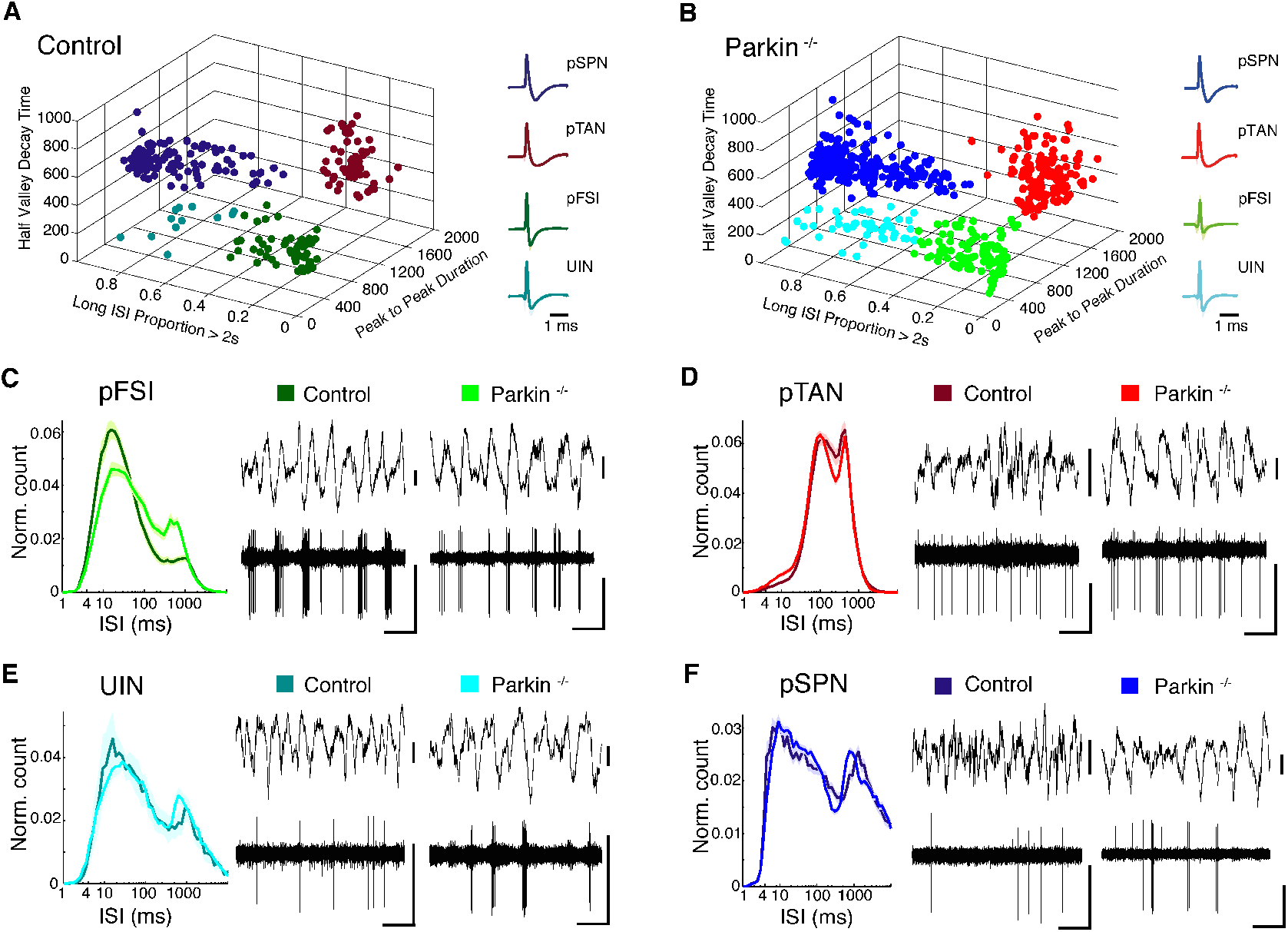
Classification of putatively different striatal neuron types in control and parkin^−/−^ mice based on electrophysiological and waveform parameters. (A, B) Left side, In both control (A) and parkin^−/−^ mice (B), multidimensional clustering reveals 4 putatively different subclasses of neurons, with each dot representing a single unit. In this and all following panels, a specific colour is assigned to each neuron type with dark colors generally representing control data and bright colors representing data from parkin mutant mice (blue: pSPN, red: pTAN, green: pFSI, cyan: UIN). Right side, Mean waveform +/− std shading for each population from control (A) and parkin^−/−^ mice (B). Note similar clustering results in A and B for both experimental groups. (C-F) Left panels, normalized mean ISI histogram for each neuronal subpopulation in control (dark) and parkin^−/−^ group (bright). +/− SEM shading on a logarithmic x-axis. pTANs exhibit a uniform distribution, reflecting their unique tonic discharge pattern. Right panels, exemplary raw data traces for each putative neuron group from controls (left) and mutants (right) with simultaneously recorded cortical LFP for pFSI (E), pTAN (D), UIN (E), pSPN (F). Horizontal scale bars: 1s, vertical scale bars: 0.05 mV. Control mice and parkin deficient mice reveal similar neuronal subclasses and share an identical data distribution. Supplementary Figure 3 reveals an identical spatial sampling for neurons recorded in the healthy and parkin mutant striatum, respectively.

Because numbers of FSIs or Parvalbumin-positive neurons express a decreasing gradient from lateral to medial in the striatum (Gerfen CR et al. 1985) we in addition analyzed the spatial distribution of depicted neuron types in different sections of the striatum shown in supplementary Figure 3.

### Parkin deficiency alters key firing properties of striatal pFSIs

To gain insight into the pathophysiological relevance of parkin mutations on neural activity within the striatal microcircuitry, we tested for changes in the ongoing discharge characteristics of each neuronal subtype. Figure 2 provides an overview on quantitative comparisons of firing rates, bursting characteristics and discharge regularity (as measured by the local coefficient of variation of ISIs, CV2) of striatal interneurons and projection neurons in controls and parkin^−/−^ mice.

**Figure 2.**
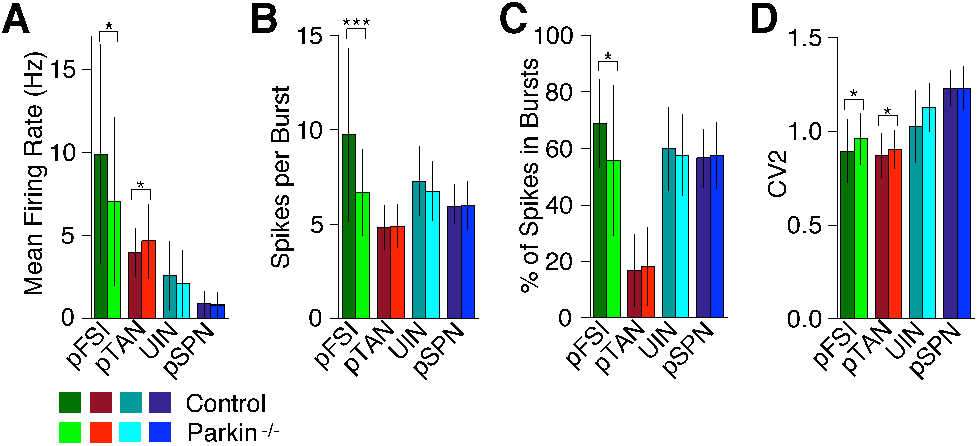
Striatal neuronal discharge properties in controls and parkin^−/−^ mice. (A-D) Bar graphs represent mean values +/− std of mean firing rate (A), spikes per burst (B), % of spikes in bursts (C) and CV2 (D) for all neuronal subpopulations. Color code as above. Pairwise comparisons revealed significant differences for the discharge rates of putative striatal interneurons (pFSIs, pTANs) but not putative projection neurons (pSPNs) after correction for multiple comparisons (A). Decreased spiking of parkin mutant pFSIs is reflected by significantly fewer spikes per burst and a reduced amount of spikes contained within bursts, respectively (B, C). Higher values for the local coefficient of variation (CV2) indicate that mutant interneuronal discharge (pFSIs/pTANs) is significantly more irregular compared to controls (D). * *p* < 0.05, *** *p* < 0.001.

Consistent with the absence of obvious motor symptoms in parkin mutant mice, discharge rates of pSPNs were similar in both experimental groups (ctrl, 0.9±0.8 Hz vs. parkin^−/−^, 0.8±0.8 Hz; MWUt [n=121/324], *p*=0.46). While firing rates of UINs were also similar in controls and parkin^−/−^ mice (ctrl, 2.6 ± 2.1 Hz vs. parkin^−/−^, 2.1 ± 2.0 Hz; MWUt [n=12/47], *p*=0.38), the presence of parkin mutations had a significant impact on the discharge characteristics of pFSIs. We found a significant reduction (∼30%) in the discharge rate of pFSIs in parkin^−/−^ mice (ctrl, 9.9 ± 6.6 Hz vs. parkin^−/−^, 7.0 ± 5.1 Hz; MWUt [n=56/117], *p*=0.017). This effect was likely driven by a reduced intra-burst spike count of mutant pFSIs (ctrl vs. parkin^−/−^, 9.8 ± 2.3 vs. 6.7 ± 1.8 spikes/burst; MWUt, *p*=3.7e^−6^) and significantly fewer spikes contained in bursts (ctrl vs. parkin^−/−^, 68.9 ± 15.6% vs. 55.8 ± 26.7%; MWUt, *p*=0.038). pTANs showed an opposing trend: Their firing rate was slightly, but significantly, elevated in mutant mice when compared to controls (ctrl, 4.0 Hz ± 1.4 Hz vs. parkin^−/−^, 4.7 Hz ± 2.2 Hz; MWUt [n=63/154], *p*=0.04). In addition to these changes in firing rate and burst structure, spike train irregularity (CV2) was also selectively increased in mutant pFSIs (control, 0.9 ± 0.2 vs. parkin^−/−^ 1.0 ± 0.1; MWUt, *p*=0.0499) and pTANs (ctrl, 0.8 ± 0.1 vs. parkin^−/−^, 0.9 ± 0.1; MWUt, *p*=0.0499), but not pSPNs (ctrl, 1.2 ± 0.1 vs. parkin^−/−^, 1.2 ± 0.1; MWUt, *p*=0.771). We conclude that, in a preclinical stage, mutations in the parkin gene are associated with excitability changes in local striatal interneuronal populations, with discharge properties of projection neurons being unaltered.

### Correlation structure of the striatal microcircuit under isoflurane anesthesia

We next sought to understand the consequences of altered interneuronal excitability on the correlation structure of the striatal microcircuitry. To this end, we assessed the coupling strength between all recorded pairs of striatal neurons by calculating rate-normalized cross-correlation functions for all neuronal subtypes. In general, striatal single cell activity was highly correlated in both groups under isoflurane anesthesia (∼70% of all cross-correlograms were significantly modulated according to the criteria described in the methods section), and displayed synchronized slow (∼1-2 Hz) oscillatory burst discharges with a marked predominance (∼90%) of positive correlations. The mean pair-wise correlation strength did not differ significantly between knockouts and controls in any of the tested groups (supplementary Table 2), with the notable exception of pFSI pairs.

### Disruption of the local striatal pFSI circuitry

In line with the tight electrotonic and electrochemial coupling described for this cell type (Kita et al., 1990; Koós and Tepper, 1999; Fukuda, 2009), neuronal activity of simultaneously recorded pFSIs in control animals was highly positively correlated. In contrast, pFSI-pFSI synchrony was generally weaker in parkin^−/−^ mice (t-score of ctrl, 29.5 ± 14.8 vs. parkin^−/−^, 15.2 ± 8.5, MWUt [n=30/31], *p*=9.1e^−5^, Figure 3A). We then partitioned pFSI pairs by distance to study their correlations as a function of intercellular distance (Figure 3C). Nearby pFSI pairs in the healthy striatum were positively correlated with high magnitude. In agreement with the intrastriatal short-range connectivity profile recently described for FSIs (Straub et al., 2016), pFSI-pFSI synchrony showed a marked distance-dependent decay (<1 mm vs. 1-2 mm, MWUt [n=7/16], *p*=0.0056; 1-2 mm vs. >2mm, MWUt [n=16/8], *p*=0.0037; Figure 3C, left panel). In contrast, local pFSI coupling in mice with parkin mutation was strongly reduced and the spatial structure of correlated FSI firing absent (<1 mm vs. 1-2 mm, MWUt [n=14/18], *p*=0.13; 1-2 mm vs. >2mm, MWUt [n=18/9], *p*=0.94; Figure 3C, right panel).

**Figure 3.**
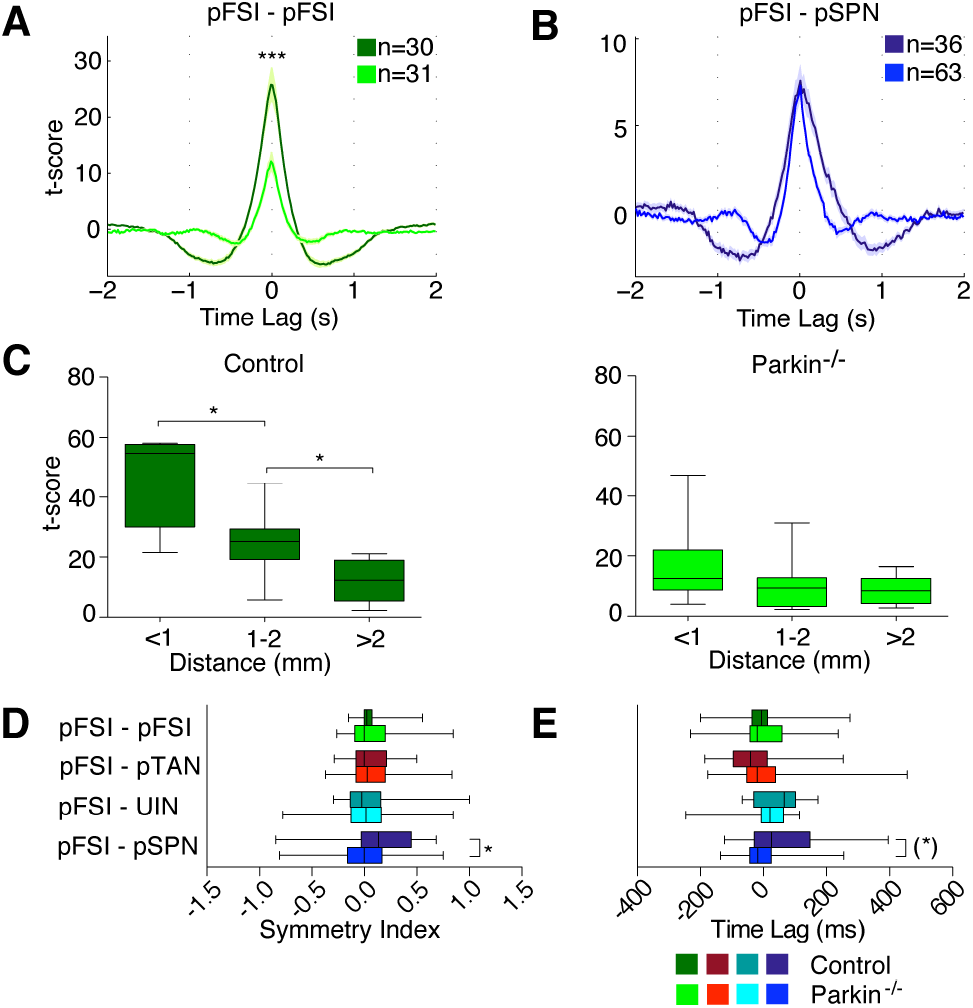
Cross-correlation analysis of pFSIs. (A) Rate normalized and t-scored cross-correlation histogram of pFSI-pFSI pairs for control animals (dark green) and parkin^−/−^ mice (bright green). Shading is SEM. Synchronization strength is significantly weaker in parkin mutant mice. (B) Rate normalized and t-scored cross-correlation histogram of pFSI-pSPN pairs for controls (dark blue) and parkin deficient mice (bright blue). Maximum t-score is not different, but note that the characteristic asymmetric shape of the central peak is absent in the parkin^−/−^ group. Cross-correlation histograms for all remaining pairs of neuronal subtypes are shown in supplementary Figure 4. (C) Mean t-score plotted as a function of distance between simultaneously recorded and cross-correlated pFSI pairs. Left panel, In controls, the average t-score diminishes progressively as a function of intercellular distance, indicating a physiological pattern of strong local and weak distant coupling among pFSIs, respectively. Right panel, This distance dependent decay of pFSI-pFSI coupling strength is absent in parkin mutant mice. (D, E) Temporal relationship of pFSI discharges with other striatal subpopulations of neurons. (D), Symmetry index of the center peak of cross-correlation histograms between pFSIs and other neuronal subclasses. Note the asymmetric distribution for pFSI-pSPN pairs in controls but not parkin mutant mice. (E) Time lag of the center peak of cross-correlation histograms between pFSIs and other neuronal subclasses. For mutant pFSI-pSPN pairs, the average time lag is shifted towards negative values. Thus, the physiological spiking sequence of pFSIs preceding pMSN discharges seen in control animals is absent in mice with a mutation in the parkin gene. (A-E) While pFSIs always serve as the reference cell, the color code in B,E and F refers to the identity of the trigger cell (pFSI-pFSI ctrl: n=36, parkin^−/−^ n=63; pFSI-pTAN ctrl: n=41, parkin^−/−^ n=46; pFSI-UIN ctrl: n=8, parkin^−/−^ n=14, pFSI-pSPN ctrl: n=36, parkin^−/−^ n=63). Whiskers of the box plots show the 5-95 percentile. (*) *p*<0.05 before correction with FDR, * *p*<0.05, \*\*\**p*<0.001 after correction. Detailed descriptive statistics for all available auto- and cross-correlations are given in supplementary Table 1 and 2.

Given the powerful inhibitory influence of bursting FSIs on striatal outflow (Friedman et al., 2017; Lee et al., 2017; O’Hare et al., 2017), globally reduced activity of and correlation levels and among FSIs should have an effect on the functional interaction between pFSIs and pSPNs. In awake non-human primates, cross-correlograms between these two cell classes are asymmetric (Adler et al., 2013), with fast activation of FSIs and delayed spiking of SPNs in response to common input. As depicted in Figure 3B, asymmetric pFSI-pSPN coupling was indeed present in healthy controls, but absent in transgenic animals. Histogram peaks were significantly shifted toward positive values in control animals—indicating trigger cell (pFSI) discharge before the reference cell (pSPN)—but symmetrically distributed around zero in mutant mice (symmetry index of ctrl, 0.18 ± 0.33 vs. parkin ^−/−^, −0.03 ± 0.33; MWUt [n=36/69], *p*=0.0215; Figure 3D). Furthermore, center peaks of pFSI-pSPN cross-correlograms were shifted towards earlier firing of pFSIs (i.e., prior to pSPNs) in the control group but not in parkin transgenic mice (time lag of ctrl, 76 ± 148 ms vs. parkin^−/−^, −2 ± 80 ms; MWUt [n=36/69], before correction: *p*=0.0091, after FDR correction: *p*=0.0816; Figure 3E). Taken together, these findings suggest that parkin deficiency leads to weaker interneuronal synchrony among striatal pFSIs and a selective timing disruption within the striatal FSI-SPN microcircuit.

### Cortico-striatal cross-correlations

Previous in vitro studies have identified the cortico-striatal synapse as an important functional locus for changes in parkin deficient mice (Kitada et al., 2009; Martella et al., 2009; Madeo et al., 2012). To examine alterations along the cortico-striatal axis as a possible mechanism behind the observed timing disruptions of the mutant pFSI microcircuitry, we calculated spike train cross-correlation functions between simultaneously recorded multiunit clusters in motor cortex and striatal single units. Significant features were detected in 590 of 648 (91%) cross-correlograms. Of these, the vast majority (544/590, 92%) had broad positive central peaks at around zero lag (Figure 4A-D), demonstrating slow oscillatory covariation of firing rates (i.e., synchronized bursting) in cortex and striatum. We noted a slight, but non-significant asymmetry of the central peak in cross-correlograms constructed from cortex-pFSIs pairs in controls compared to knockouts (symmetry index of ctrl, 0.02 ± 0.14 vs. parkin^−/−^, −0.07 ± 0.29; MWUt [n=49/102], before correction *p*=0.03, FDR-corrected *p*=0.112; Figure 4A,E). However, we observed a significant lead of cortical multiunits over both pSPNs and pFSIs in parkin mutant mice, but not in healthy controls (time lag of Cx-pSPN pairs in ctrl, 15 ± 81 ms vs. parkin^−/−^, −32 ± 77ms; MWUt [n=56/165], *p*=0.0010; Figure 4B,F; time lag of Cx-pFSI pairs in ctrl, −6 ± 72 ms vs. parkin^−/−^, −39 ± 78 ms; MWUt [n=49/102], *p*=0.0019; Figure 4A,F).

**Figure 4.**
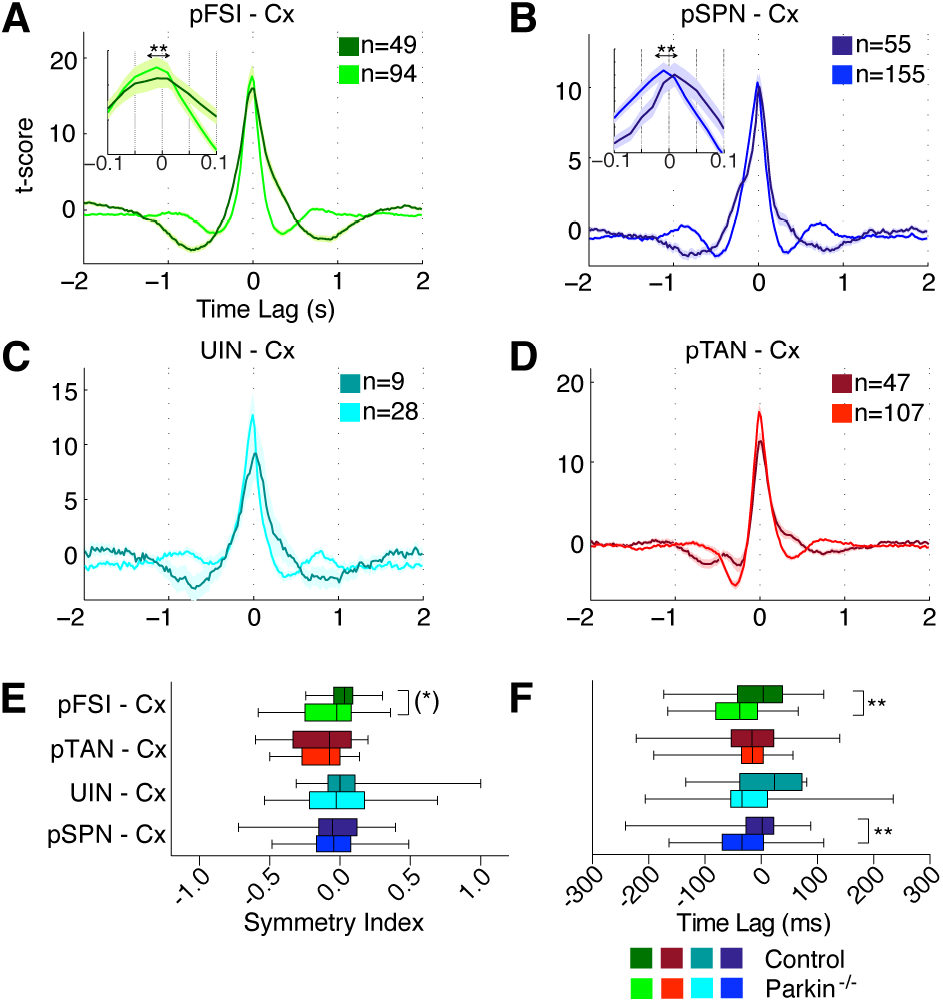
Cross-correlations between striatal neurons and cortical multiunits. (A-D) T-scored cross-correlation histograms constructed from pairs of positively correlated striatal single units and multiunits recorded in motor cortex. Insets in A and B highlight the significant time shift difference of pFSI-Cx and pSPN-Cx pairs between the experimental groups. Note that negatively correlated pairs are shown in supplementary Figure 4. Further descriptive statistics are given in supplementary Table 2. (E, F) Temporal relationship between motor cortical multiunits and different striatal cell types. (E) The mean symmetry index for pFSI-Cx pairs is shifted towards negative values in the parkin^−/−^ group. Note that this difference is statistically signifcant only before correction for multiple comparisons. (F) Compared to controls, center peak time lags of cross-correlogram histograms derived from pairs of mutant motor cortical multiunits and pFSIs and pSPNs are shifted significantly towards negative values. (A-F) Color code as in previous figures with striatal cells as reference. Brighter and darker colors refer to group data from parkin mutants and healthy controls, respectively. Whiskers of the box plots show the 5-95 percentile. (*) *p*<0.05 before correction with FDR, ** *p*<0.01.

### Unit cross-correlations along the cortico-striato-pallidal axis

Given these intrastriatal microcircuit changes in parkin mutant mice, we next sought to determine consequences of these alterations on downstream activity in the LGP. Compared to controls, the average discharge rates of single LGP units was slightly, but not significantly, elevated in the parkin^−/−^ group (ctrl, 24 ± 16 Hz; parkin^−/−^, 31 ± 20 Hz; MWUt [n=55/55], before correction *p*=0.044, FDR-corrected *p*=0.07) (Figure 5). Likewise, neither bursting behaviour nor discharge regularity differed between the two experimental groups (percentage of spikes in bursts, ctrl, 18.9 ± 14.6% vs. parkin^−/−^, 16.8 ± 15.1%; MWUt [n=54/53], *p*=0.56; CV2, ctrl, 0.6 ± 0.2 vs. parkin^−/−^, 0.6 ± 0.2; MWUt [n=55/55], *p*=0.39). In agreement with previous observations (Goldberg et al., 2003; Magill et al., 2006; Walters et al., 2007; Zold et al., 2007; Mallet et al., 2008, 2012; Abdi et al., 2015; Karain et al., 2015), rhythmic LGP bursting was tightly coupled to both cortical and striatal discharges at slow-wave frequencies, indicating a highly synchronized, oscillating cortico-striato-pallidal circuit (Figure 5).

77 out of 97 (81.3%) Cx-LGP pairs were significantly modulated with approximately equal proportions of positive and negative correlations (resulting from pallidal neurons exhibiting in-phase and anti-phase spiking with cortical units, respectively) in both experimental groups (Figures 5 and 6, supplementary Table 2). It is of note that pallidal neurons also exhibited slow oscillatory in-phase and anti-phase synchrony with both putative striatal projection neurons (Figure 6C) and interneuron populations (Fig. 6B and D). The timing of pallido-striatal spiking as assessed by cross-correlation analysis did not differ between the experimental groups (Figure 6, E and F). The point to be noted from the symmetry-analysis depicted in Figure 6E is that, in both groups, cross-correlogram peaks constructed from pairs of pallidal neurons and striatal interneurons (pFSI/pTAN) were generally shifted towards negative values, while central peaks derived from cortico-pallidal and SPN-pallidal pairs were shifted towards positive values.

**Figure 5.**
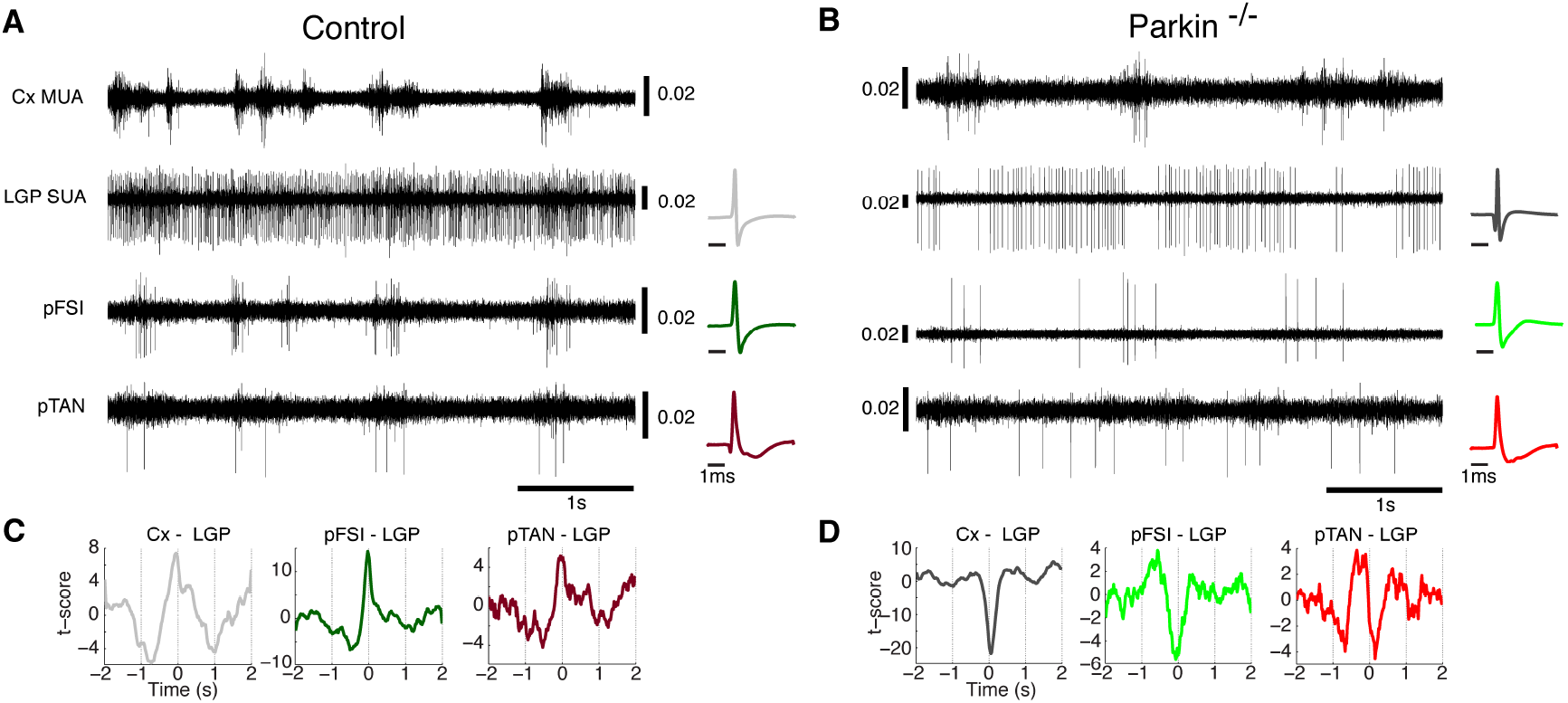
Illustrative examples exhibiting different coupling modes of unit activities along the cortico-striato-pallidal axis during isoflurane anesthesia. Raw traces are taken from a recording in a control (A) and a parkin mutant animal (B), respectively. In both panels, a cortical multiunit (Cx MUA), is depicted together with pallidal single units (LGP SUA) and two types of striatal interneurons (pFSI and pTAN). Vertical scale bars for the individual units are given in mV. Mean action potential waveforms of the unit are shown on the right. Positive and negative rate covariations are readily discernible. (C, D) Cross-correlation histograms of the units shown in (A) and (B). Both positive and negative coupling patterns were similarly observed in both experimental groups (see results).

**Figure 6.**
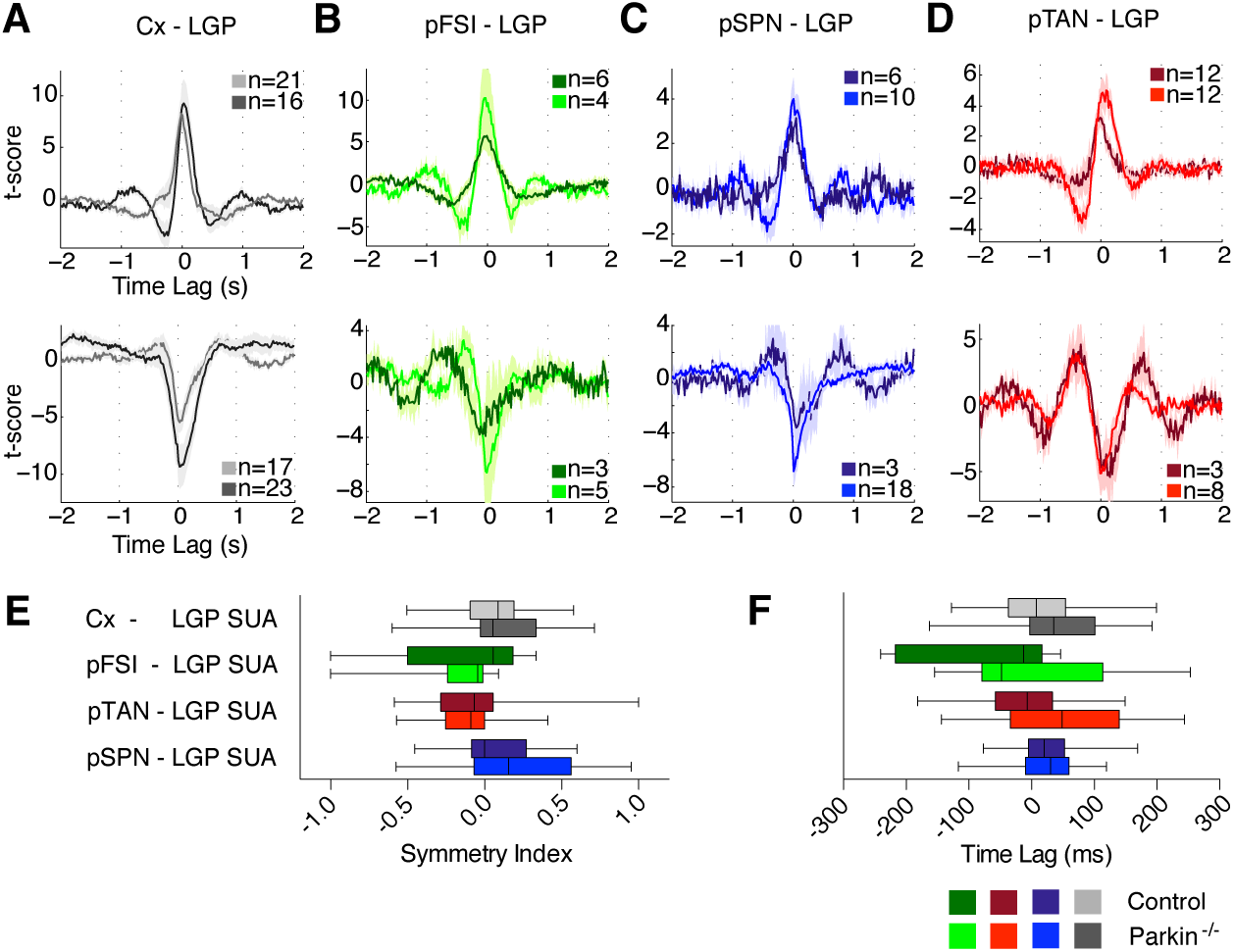
Temporal relationships between pallidal single units, different striatal neuron types and motor cortical multiunits. (A-D) Mean t-scored cross-correlogram histograms with SEM shading for all pairs of Cx-LGP (A), pFSI-LGP (B), pTAN-LGP (C) and pSPN-LGP (D). Upper and lower panels show group averages for positively and negatively correlated pairs, respectively. Note similar proportions of phase and anti-phase correlated Cx-LGP pairs in both groups. In all panels, spiking of LGP single units is triggered on cortical and striatal unit activities, respectively. (E) Symmetry index for center peaks of all cross-correlation histograms shown in A-D. In both experimental groups, symmetry indices derived from cross-correlated pairs of pallidal neurons and pSPNs are positive, whereas those derived from cross-correlated LGP units and striatal interneurons are shifted towards negative values. Thus, pallidal spiking generally lagged striatal outflow activity but preceded striatal interneuronal discharges. Similar temporal relationships can be seen in the lag analysis of cross-correlation center peaks (F). Colour code as noted above. Whiskers of box plots show 5-95 percentile. For detailed descriptive statistics, the reader is referred to supplementary Table 2.

Thus, under conditions of isoflurane anesthesia, spiking of cortex and SPNs preceeds LGP units, the latter at the same time leading spiking of striatal interneurons. While the overall proportion of significantly modulated striato-pallidal cross-correlograms was not different between the two groups (ctrl, 33 out of 89 (37%) pallido-striatal CCFs vs. parkin^−/−^, 57 out of 117 (49%), Fisheŕs exact test, *p*=0.12) we noticed a higher prevalence of negative pallido-striatal cross-correlations among the group of parkin mutant mice (ctrl, 9 out of 33 pairs (27%) vs. parkin^−/−^, 31 out of 57 (54%); Fisheŕs exact test, *p*=0.016). Thus, under general anesthesia with isoflurane, a large-scale cortex-basal ganglia network is highly synchronized and tuned to the prevalent slow wave rhythm of the cortex. While knockout animals exhibited cell-specific abnormalities within intrastriatal interneuronal microcircuits, spiking activity along the striato-pallidal pathway (pSPNs to LGP) was not altered in parkin mutant mice.

### Parkin-deficiency is associated with amplified beta oscillations in striatal and pallidal LFPs

We next asked whether parkin deficiency might also alter global population network dynamics (as reflected in local field potentials (LFPs)) within and across the corresponding regions. Figures 7 A and B show representative time-frequency transformed records of ongoing LFPs measured simultaneously in Cx, CPu and LGP of a control and a parkin mutant mouse, respectively.

**Figure 7.**
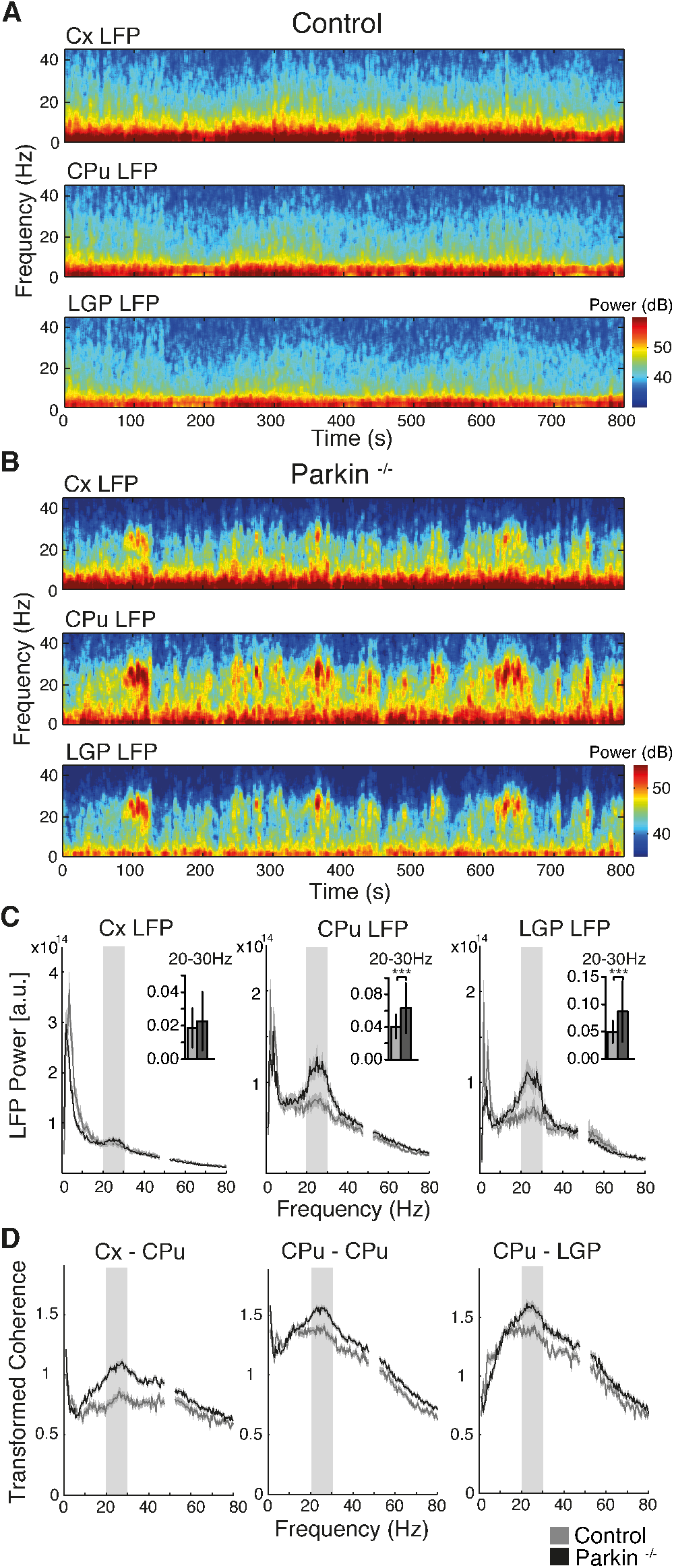
Local field potentials reveal enhanced synchronized beta oscillations in striato-pallidal circuits of parkin mutant mice. (A,B) Example time-frequency transformed LFP power of simultaneous recordings from motor cortex, CPu and LGP from a control (A) and parkin^−/−^ mouse (B). (C) Mean relative 1/f normalized group power spectra of LFP recordings in motor cortex (Cx, left, ctrl: n=84, parkin^−/−^: n=190), CPu (middle, ctrl: n=86, parkin^−/−^: n=196) and LGP (right, ctrl: n=56, parkin^−/−^: n=87) +/− SEM shading of control (grey) and parkin^−/−^ mutant mice (black). Insets depict relative power in the beta-frequency range (20-30 Hz; mean +/− std). Note that beta power is significantly elevated in CPu and LGP, but not motor cortex, of parkin deficient animals. (D) Transformed group coherence (+/− SEM) between LFP pairs (Cx-CPu (left) ctrl: n=79, parkin^−/−^: n=180, CPu-CPu (middle) ctrl: n=81, parkin^−/−^: n=185 and CPu-LGP (right) ctrl: n=61, parkin^−/−^: n=97). Beta band coherence is significantly enhanced along the entire mutant cortico-striato-pallidal axis. \*\*\**p*<0.001.

In both experimental groups, LFP power spectral density was dominated by a large peak in the slow-wave frequency band in all three structures (Figure 7C). Importantly, global population dynamics at slow wave frequencies did not differ between knockouts and controls at a cortical level, neither for absolute (MWUt, [n=84/190], *p*=0.05) nor relative (MWUt, [n=84/190], *p*=0.06) power measures. Unexpectedly, and in contrast to the control group, a prominent second peak emerged in ongoing striatal and pallidal, but not cortical, LFPs of parkin mutant mice in the beta frequency band (mean peak frequency, 23 ± 3 Hz; Figure 7C). A quantitative comparison revealed that motor cortex LFP power in the beta-frequency band was similar (relative power between 20 and 30 Hz, MWUt [n=84/190], *p*=0.28) or even reduced (absolute beta power; MWUt [n=84/190], *p*=0.02) in parkin^−/−^ mutant mice compared to controls. In contrast, a significant beta peak was consistently (6 out of 7 animals) found in the averaged absolute and relative power spectra LFPs recorded from CPu and LGP of parkin deficient mice, respectively (comparison of absolute beta power for CPu of ctrl vs. parkin^−/−^; MWUt [n=86/196], *p*=1.36e^−07^; LGP, MWUt [n=56/87], *p*=6.59e^−05^; comparison of relative beta power for CPu of ctrl vs. parkin^−/−^; MWUt [n=86/196], *p*=2.13e^−09^; LGP, MWUt [n=56/87], *p*=0.0001; Figure 7C). The spectrograms depicted in Figure 7B demonstrate the waxing and waning dynamics of prominent ongoing beta oscillatory episodes, which emerge synchronously along the cortico-striato-palllidal route of parkin mutant mice. Their burst-like appearance is reminiscent of the fine temporal structure of exaggerated beta oscillations within the human parkinsonian basal ganglia circuit (Moll et al., 2015; Tinkhauser et al., 2017; Sharott et al., 2018). Cortico-striato-pallidal network synchronization was next assessed by computing coherence spectra of simultaneously recorded LFPs within and across the recorded structures. Mean transformed coherence in the slow wave frequency range did not differ between the experimental groups along the cortico-striatal axis (Cx-CPu MWUt [n=79/180], *p*=0.29 and CPu-CPu pairs MWUt [n=81/185], *p*=0.62) but showed decreased coherence for CPu-LGP (MWUt [n=61/97], *p*=2.87e^−05^) and Cx-LGP (MWUt [n=57/78], *p*=0.0006, (not shown)) pairs in the parkin^−/−^ group. Figure 7D demonstrates that intra- and interstructural coherence spectra of knockouts peaked at beta frequencies (peak frequency, 25 ± 2Hz), with significantly stronger synchrony in the beta frequency range compared to controls (comparison of beta-band (20-30Hz) LFP coherence between ctrl and parkin^−/−^; Cx-CPu, MWUt [n=79/180], *p*=9.55e^−13^; CPu-CPu, MWUt [n=81/185], *p*= 5.83e^−06^; CPu-LGP, MWUt [n=61/97], *p*=7.93e^−06^; Cx-LGP (not shown), MWUt [n=57/78], *p*=7.88e^−12^). Together, these data show that—at the LFP level—parkin deficiency leads to the emergence of amplified beta oscillations in striato-pallidal circuits that are coherent with cortex, resembling a cardinal electrophysiological signature of chronic dopaminergic denervation in toxin-based animal models (Sharott A et al. 2005; Mallet N, A Pogosyan, LF Marton, et al. 2008; Mallet N, A Pogosyan, A Sharott, et al. 2008; Avila I et al. 2010; Brazhnik E et al. 2012; Lemaire N et al. 2012; Beck MH et al. 2016) and patients with Parkinsońs disease (Brown P et al. 2001; Levy R, P Ashby, et al. 2002; Levy R, WD Hutchison, et al. 2002; Williams D et al. 2002; Kuhn AA et al. 2008; Bronte-Stewart H et al. 2009; Moshel S et al. 2013; Sharott A et al. 2014; Moll CK et al. 2015; Sharott A et al. 2018).

### Parkin deficiency perturbs striatal spike-LFP relationships and leads to amplified coupling of pFSIs to ongoing beta oscillations

Given that parkin deficiency produced an electrophysiological phenocopy of the dopamine-denervated cortico-basal ganglia network, we finally sought to examine how amplified population synchrony might reshape the phase coupling of spikes to LFP oscillations. Based on the results of the power spectral density analysis, we focused our analysis on two dominant frequency bands: slow wave frequency (1.5-3.5 Hz) and beta frequency (20-30 Hz). In line with previous observations (Sharott et al., 2009, 2012), the vast majority of striatal neurons (∼80%) in both genotypes exhibited significant phase coupling of their spikes to cortical slow waves. However, parkin mutations were generally associated with a significantly strengthened spike coupling to ongoing cortical slow wave oscillations, irrespective of striatal cell type (comparison ctrl vs. parkin^−/−^; pFSI, MWUt, [n=48/96], *p*=6.57e-06; pTAN MWUt, [n=57/125], *p*=0.001; UIN, MWUt, [n=9/28], *p*=0.006; pSPN, MWUt, [n=84/249], *p*=0.004; Figure 8A). pFSI and pSPN spiking in the parkin knockout occurred at a later phase of the cortical LFP compared to controls, in keeping with the results of the cortico-striatal unit cross-correlation analysis (comparison ctrl vs. parkin^−/−^; pFSI, WWFt [*F*(1,142)=8.41], *p*=0.01; pSPN, WWFt [*F*(1,331)=23.21], *p*=1.11e^−05^; Figure 8A). In accord with observations in a toxin-based animal model of PD (Lemaire et al., 2012), the intrastriatal phase preference of all striatal subtypes of neurons was significantly shifted in knockout animals (comparison ctrl vs. parkin^−/−^; pFSI, WWFt F(1,137)=4.65, *p*=0.03; pTAN, WWFt F(1,184)=10.23, p=0.004; UIN, WWFt, F(1,48)=7.73, *p*=0.0097; pSPN, F(1,340)=7.94, *p*=0.0085; Figure 8B). Interestingly, coupling of mutant pSPNs and pFSIs to pallidal LFPs was also significantly enhanced, although this was not significant for pFSIs after correction for multiple comparisons (comparison of ctrl vs. parkin^−/−^, pSPN, MWUt [n=41/114], *p*=0.0089; pFSI, MWUt [n=19/39], *p*=0.0276 before correction, FDR-corrected *p*=0.7; Figure 8C). Figure 8C also demonstrates that—in contrast to the forward shift in the preferred phase in relation to cortical and striatal LFPs observed in knockouts, respectively—spikes of mutant pSPNs and pFSIs (as well as pTANs) were shifted backward (i.e., occurred at an earlier phase of the pallidal LFP). Overall, genotype had no effects on phase coupling of pallidal spikes to cortical, striatal and pallidal slow waves, neither on locking-strength (MWUt, all *p* >0.05) nor phase preference (WWFt, all *p* >0.05).

**Figure 8.**
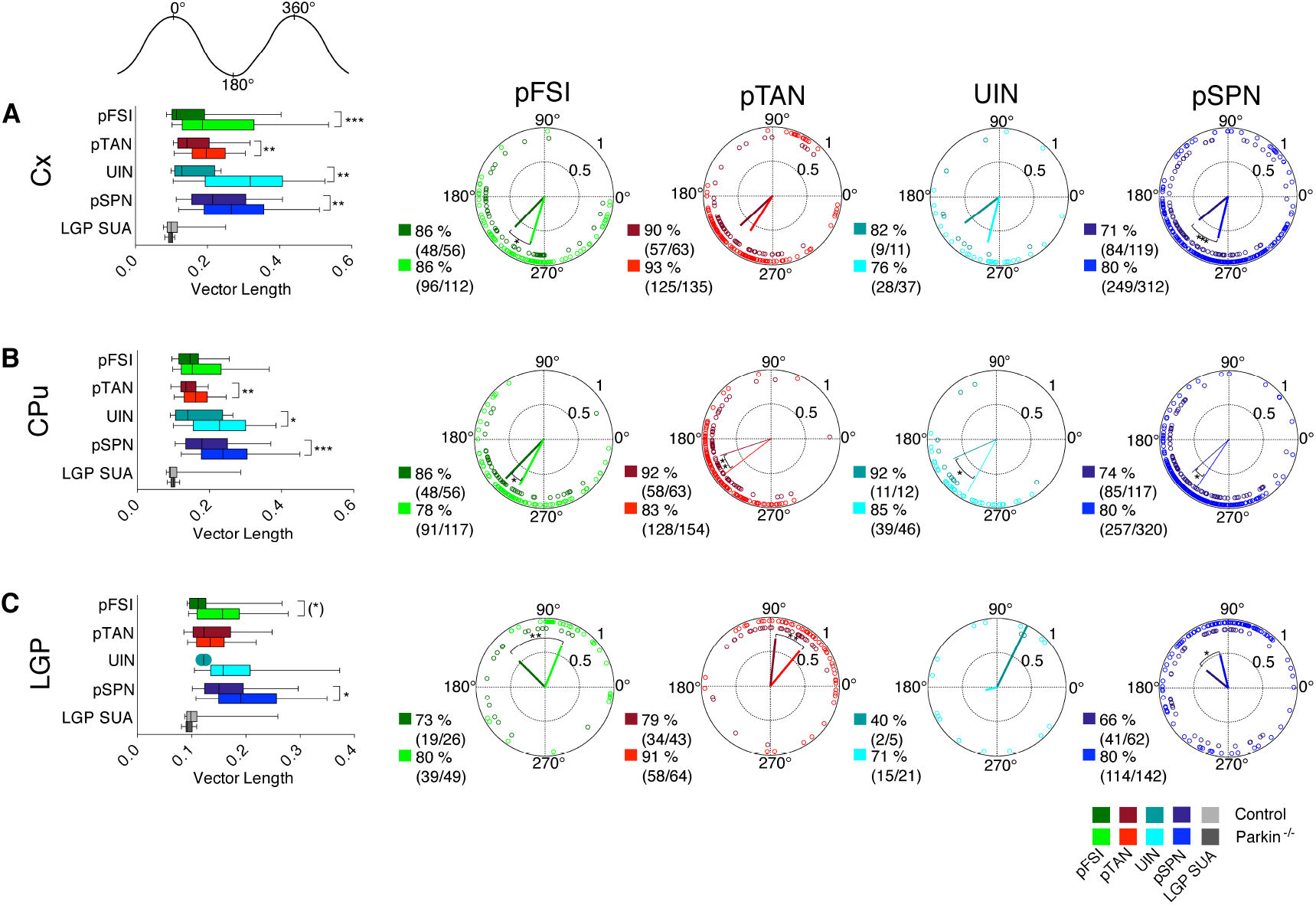
Low-frequency phase locking of units to LFP. In both experimental groups, all striatal neuron subclasses of neurons and LGP single units were strongly and significantly phase locked to the cortical (A, top row), striatal (B, middle row) and pallidal (C, bottom row) LFPs filtered in the low frequency range of slow wave activity dominating under isoflurane anesthesia (i.e. between 1.5-3.5 Hz). (A-C), Left panels, pairwise comparisons of the vector length (a rate-normalized indicator of phase locking strength) for all neuron types and neurons that were significantly phase-locked to LFP activities, respectively. Note the significant stronger phase locking of all striatal neurons to cortical LFPs in mice with parkin deficiency. A subset of mutant striatal neuron types (pSPNs, UIN and pTANs) were also stronger locked to striatal LFPs compared to controls. Spike-field-locking of main striatal outflow (pSPNs) to pallidal LFPs was significant higher for parkin deficient mice, while it was for pFSI only significant before correction for multiple comparisons. The circular plots in right panels show mean phase angles for neurons that were significantly locked to LFPs from motor cortex (A), CPu (B) and LGP (C), respectively. A dot represents the mean phase angle of an individual neuron. Vector length is derived from the variance of all mean angles. A phase angle of 180° corresponds to the negative peak of the LFP oscillation. For each group, both percentage and absolute number of significantly locked neurons are stated. Note the significant phase shift of almost all striatal neuron types with respect to the negative peak of ongoing slow wave LFP oscillations in the parkin^−/−^ group. As above, darker and brighter colors in all panels represent data from controls and parkin^−/−^ mice, respectively. Whiskers of box plots denote the 5-95 percentile. (*) p<0.05 before correction, *p<0.05, ** p<0.01, *** p<0.001.

Finally, our finding of exaggerated beta oscillations in striatal and pallidal LFPs of parkin mutant mice raised the possibility that these rhythms could also be represented in the spike trains of individual striatal or pallidal neurons. Phase-locking strength of all subtypes of striatal neurons, however, was not different between the two groups (MWUt, all *p* >0.05; supplementary Figure 5). Notably, however, we observed an almost doubled proportion of pFSIs that were significantly phase-locked to cortical and striatal beta-oscillations in the parkin^−/−^ group—although the statistics were significant only before correction for multiple comparisons (ctrl, 10 out of 56 pFSI/Cx-LFP pairs (18%) vs. parkin^−/−^, 40 out of 112 (36%); Fisheŕs exact test, *p*=0.02; FDR-corrected *p=0.08*; ctrl, 12 out of 56 pFSI/CPu-LFP pairs (21%) vs. parkin^−/−^, 46 out of 117 (39%); Fisheŕs exact test, *p*=0.03; FDR-corrected *p*=0.1; Figure 9A-C).

**Figure 9.**
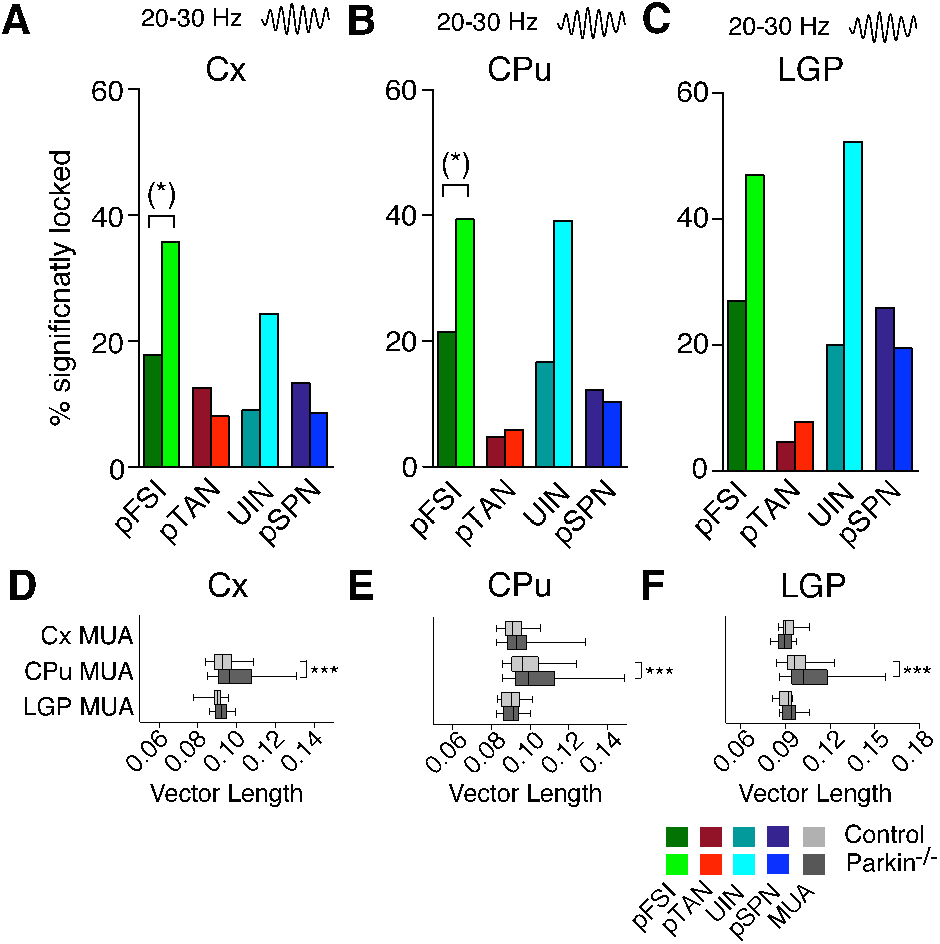
Phase locking of unit activities to LFP oscillations in the beta frequency range (20-30Hz). (A-C), Percentage of significantly locked striatal neuronal subpopulations to bandpass-filtered (20-30Hz) LFPs recorded in motor cortex (A), CPu (B) and LGP (C). Note that the proportion of pFSIs and UINs significantly locked to beta oscillations is nearly doubled in the parkin^−/−^ group. However, the higher prevalence of beta-locked pFSIs in mutants compared to controls reached significance only before correction for multiple comparisons. The corresponding bar graphs of vector length and circular plots are shown in supplementary Figure 5. (D-F), Mean vector length for phase locking of Cx-MUA, CPu-MUA and LGP-MUA to cortical (D), striatal (E) and pallidal beta oscillations (F). Note that the phase locking strength of CPu-MUA to all available LFPs is selectively enhanced. Whiskers of boxplots show 5-95 percentile. (*) *p*<0.05 before FDR-correction, \*\*\**p*<0.001

Compared to single unit spike trains, recordings from local clusters of neighbouring neurons around the electrode tip may be more susceptible to changes in high-frequency oscillations (Fries P et al. 2001) and coherence with the LFP (Zeitler M et al. 2006). We therefore conducted the same set of analyses as above for multiunit signals recorded from the corresponding structures. Despite the lack of strong effects of genotype on the strength of striatal single-unit entrainment to beta oscillations, parkin mutations were associated with a significantly stronger phase-locking of striatal *multiunits* to beta-band oscillations recorded from cortex, CPu and LGP (CPu-MUA/Cx-LFP; MWUt [n=196/531], *p*=6.26e^−08^; CPu-MUA/CPu-LFP; MWUt [n=259/671], *p*=2.36e^−05^; CPu-MUA/LGP-LFP; MWUt [n=151/301], *p*=3.44e^−06^; Figure 9D-F). This was accompanied by a significant increase in the percentage of striatal multiunits significantly phase-locked to beta-band oscillations of LFPs from all three structures (Fisheŕs exact test, all *p* <0.05, exact values are shown in supplementary Table 3). Notably, neither coupling prevalence (see supplementary Table 3) nor coupling strength (Cx-MUA/CPu-LFP; MWUt [n=55/104], *p*=0.13; Cx-MUA/LGP-LFP; MWUt [19/34], *p*=0.20; LGP-MUA/Cx-LFP; MWUt [n=10/22], *p*=0.17; LGP-MUA/LGP-LFP; MWUt [n=5/9], *p*=0.62) of cortical and pallidal multiunits to beta oscillatory activities in the LFP was different between controls and knockouts in any structure.

## Discussion

Here, we characterized the striato-pallidal circuit in the parkin deficient mouse model in vivo. We show that—in the absence of overt nigral degeneration—mutations in the parkin gene lead to a re-arrangement of striatal microcircuits and predispose to an electrophysiological phenotype characteristic of advanced PD. Our results suggest that exaggerated oscillatory activities in the beta frequency band (20-30Hz) in striato-pallidal circuits may precede cell loss of dopaminergic neurons in the substantia nigra and may thus emerge earlier in the disease process than previously thought - which has implications for pathophysiological concepts and biomarker development.

The two major findings of this study describe adaptive changes in cortico-striato-pallidal circuits preceding nigro-striatal cell loss and phenotypic alterations. First, parkin deficiency lead to a distinct re-arrangement of the striatal interneuronal circuitry with reduced firing dynamics of FSIs. Second, the presence of parkin mutations predisposes to a focal electrophysiological phenocopy of advanced PD, restricted to striato-pallidal circuits. To the best of our knowledge, this is the first demonstration of exaggerated beta oscillations in a genetic mouse model of premanifest PD.

### Parkin deficiency leads to disruption of the striatal microcircuit

Among the different striatal circuit elements investigated in this study, the population of striatal FSI stands out prominently. Parvalbumin-containing FSIs comprise only ≈1% of all neurons in the rodent striatum (Luk KC and AF Sadikot 2001) but form a key population of striatal interneurons exerting a potent inhibitory influence upon striatal outflow (Koos T and JM Tepper 2002; Mallet N et al. 2005; Planert H et al. 2010). Moreover, the FSI circuitry is subject to powerful and convergent input from cortex (Parthasarathy HB and AM Graybiel 1997; Ramanathan S et al. 2002). The correlational structure of cortico-striatal input, in turn, is the critical driver and organizer of activity within the striatal microcircuit (Hjorth J et al. 2009; Sharott A et al. 2009). A deficit of FSI-mediated feed-forward inhibition will therefore alter the integration of cortical inputs within the striatal microcircuit. Previous research has shown that acute blockade of dopamine receptors reduces ongoing discharge rates of striatal FSIs (Wiltschko AB et al. 2010; Yael D et al. 2013), but has no effect on FSI connectivity (Gittis AH, GB Hang, et al. 2011)—indicating that the aberrant FSI spiking dynamics observed in our study may not only be due to altered dopamine receptor signaling but also to local circuit rearrangements. In this context, it is of note that dopamine depletion shifts the connectivity pattern (Salin P et al. 2009; Gittis AH, GB Hang, et al. 2011) and alters task-related ensemble activity of striatal FSIs (Hernandez LF et al. 2013). Furthermore, sub-acute remodeling of striatal FSI circuitry has been demonstrated following dopamine depletion (Gittis AH, GB Hang, et al. 2011) and presynaptic properties of GABAergic interneurons are changed (Barroso-Flores J et al. 2015). The observed reduction of FSI spiking may reflect increased gap junctional coupling (Damodaran S et al. 2014) although we observed a disruption of FSI-FSI coupling in transgenic mice suggesting that correlated inhibition at the striatal level is reduced. The spatial organization of FSI-FSI correlations as a steep function of intercellular distance in controls was repealed in parkin deficiency indicating a loss of spatial specificity. Furthermore, the asymmetric profile of FSI-SPN coupling was absent in transgenic animals, suggesting SPNs escape from FSI-mediated feed forward inhibition. In this context, the increased discharge rate of transgenic TANs may play a role since bath application of acetylcholine has been shown to reduce FSI-SPN conductance (Koos T and JM Tepper 2002). Modeling studies have suggested that the level of inter-FSI coupling determines both the overall level of striatal spiking (Hjorth J et al. 2009) and the balance between direct and indirect pathway activity (Damodaran S et al. 2014). Under physiological conditions FSIs show the fastest responses to cortical stimulation compared to other striatal circuit elements (Mallet N et al. 2005; Sharott A et al. 2012) and exert a stronger inhibitory influence on direct pathway SPNs compared to indirect pathway SPNs (Gittis AH et al. 2010). In parkin^−/−^ mice, we noted an absence of the physiological delay between pSPN with respect to pFSI spiking described in the healthy striatal microcircuit (Sharott A et al. 2009; Adler A et al. 2013). However, precise spike timing between cortex and striatum may be of ultimate importance to the regulation of cortico-striatal synaptic plasticity (Fino E and L Venance 2010). In keeping with the notion of impaired cortico-striatal plasticity observed in slices of parkin^−/−^ mice (Kitada T et al. 2009; Martella G et al. 2009; Madeo G et al. 2012), our results provide first in vivo evidence for altered cortical-striatal transmission due to parkin deficiency. Finally, reduced FSI activity in premanifest parkin^−/−^ mice may be of clinical significance since focal dystonia is a common clinical sign in parkin-associated PD preceding the onset of parkinsonian symptoms (Elia AE et al. 2014) and selective inhibition of striatal FSIs in mice has been demonstrated to evoke dystonic posturing (Gittis AH, DK Leventhal, et al. 2011).

Elevated striatal levels of acetylcholine in PD (DeBoer P et al. 1993; Ikarashi Y et al. 1997), symptom improvement through anticholinergic medication (Pisani A et al. 2007) and the reversion of hypokinetic motor symptoms by inhibition of striatal cholinergic interneurons (Ztaou S et al. 2016) highlight the critical importance of the striatal cholinergic tone in PD. Our finding of elevated activity levels of TANs (presumed cholinergic interneurons) complements recent evidence from toxin-based models (Prosperetti C et al. 2013; Chen H et al. 2018) and extend this notion to the presymptomatic disease stage in a genetic model of parkinsonism.

Motor deficits in Parkinson’s disease are commonly related to over-activity along the indirect pathway (Albin RL et al. 1989; Wichmann T and MR DeLong 1996; Kravitz AV et al. 2010). In contrast to toxin-based rodent models of PD (Tseng KY et al. 2001; Mallet N, A Pogosyan, LF Marton, et al. 2008; Kita H and T Kita 2011; Sharott A et al. 2017)— neither striato- nor pallidofugal neuronal output was significantly altered in our study. While we were not able to discriminate direct and indirect pathway SPNs, the unaltered net striato-pallidal outflow may provide a parsimonious explanation for the absence of motor signs.

On the other hand, it is well established that dopamine depletion leads to enhanced synchrony in large-scale cortico-basal ganglia networks (Tseng KY et al. 2001; Burkhardt JM et al. 2007). Consistent with previous reports in 6-OHDA depleted rats (Lemaire N et al. 2012), all striatal neuronal elements (including pSPNs) displayed increased phase coupling and phase-shifted phase locking to delta oscillations in cortical and striatal LFPs, respectively. In contrast to the described changes along the cortico-striatal axis, we found no evidence for enhanced circuit-wide synchrony of parkin-mutant neurons in the GP. This stands in contrast to the circuit-wide network synchronization of pallidal activities described in toxin-based dopamine depletion models (Raz A et al. 2001; Zold CL, B Ballion, et al. 2007; Zold CL, C Larramendy, et al. 2007; Mallet N et al. 2012; Karain B et al. 2015). Mechanistically, this may also support an important role of decorrelated GPe activity for preventing the dissemination of enhanced synchronous rhythmic activities along the entire cortico-basal ganglia-thalamo-cortical circuit (Nini A et al. 1995; Heimer G et al. 2002; Bar-Gad I et al. 2003; Edgerton JR and D Jaeger 2011).

### Enhanced synchronized striato-pallidal beta oscillations in parkin deficient mice

Enhanced beta oscillatory activity and synchronization along the cortico-basal ganglia loop have been consistently implicated to PD in humans (Brown P et al. 2001; Kuhn AA et al. 2005; Bronte-Stewart H et al. 2009; Sharott A et al. 2014) as well as toxin-based animal models (Bergman H et al. 1994; Sharott A et al. 2005; Mallet N, A Pogosyan, A Sharott, et al. 2008; Avila I et al. 2010; Tachibana Y et al. 2011; Brazhnik E et al. 2012).

The finding of exaggerated beta oscillations in the striatum and GP of parkin-deficient mice was unexpected. Both subcortical recordings in patients with advanced PD and previous work in toxin-based animal models of PD have linked the emergence of synchronized beta oscillations to late stages of severe dopamine depletion and presence of parkinsonian motor impairment, respectively (Leblois A et al. 2007; Quiroga-Varela A et al. 2013). However, the results from our study tentatively suggest that pathologically enhanced oscillatory activity in BG loops may arise earlier than previously thought. It is conceivable that parkin mutations trigger hypersynchrony in the beta frequency range focally triggered in striato-pallidal circuits by mutations in the parkin gene prior to the onset of overt neurodegeneration and the development of parkinsonian motor alterations, respectively. This finding supports the idea that early synaptic and plastic rearrangement of striatal neuronal circuits may be involved in the generation of these oscillations (Mallet N, A Pogosyan, A Sharott, et al. 2008; Degos B et al. 2009). Thus, not only extent of dopamine deficiency (as measured by overt degeneration of dopaminergic neurons), but also slow and protracted predegenerative functional disintegration of the nigro-striatal circuit may be of fundamental importance for the emergence of what could be called the prototypical electrophysiological phenotype of PD.

Interestingly, elevated beta power was detected in CPu and LGP, but not motor cortex, although cortico-striatal and cortico-pallidal coherence in this particular frequency range was enhanced. A possible explanation may be a more efficient transmission of cortical activity to downstream basal ganglia nuclei in conditions with low dopamine, leading to increased coherence, but not necessarily to elevated power at a cortical level (Brazhnik E et al. 2012). On the other hand, a striatal origin of beta oscillatory activity with lack of permeation through the basal ganglia loop to cortex is also conceivable. This concept is supported by a study showing that pharmacological cholinergic manipulation in the striatum led to the spatially confined emergence of beta oscillatory activity within the striatum—but sparing cortex (McCarthy MM et al. 2011; Kondabolu K et al. 2016). Moreover, the model of McCarthy and colleagues predicts that indirect pathway SPNs may generate rhythmicity, in line with recent findings in vivo (Sharott A et al. 2017).

### Conclusions

In summary, we described a distinct electrophysiological intra-striatal circuit reorganization together with striato-pallidal beta hypersynchrony triggered by mutation in the parkin gene. This genetically driven striato-pallidal circuit dysfunction partly resembled the electrophysiological phenotype of advanced PD and may thus reflect an early PD disease stage characterized by structurally intact but functionally compromised nigro-striatal dopaminergic circuits.

Complementary to acute toxin-based animal models of PD, genetic PD models allow for a perspective shift towards early and long-term predegenerative adaptation and compensation, as multifocal network changes develop over extended periods of time. We suggest that parkin^−/−^ mice may be a suitable model to further investigate origin and spatio-temporal evolution of pathological oscillations in premanifest stages of PD.

## Acknowledgements

This work was supported by a grant of the German Research Council (SFB 936, projects A2/A3 and C8 to A.K.E. and C.K.E.M., respectively; KR3529/4-1 to E.R. K.) and the town of Hamburg (Lexi to E.R.K.). The authors would like to thank Dorrit Bystron for her excellent technical support, Doris Lange for preparation of histological specimen, Prof. Alexis Brice and Dr. Olga Cortis for providing parkin knockout mice and Dr. Edgar Galindo-Leon for advice on data analysis.

## Supplemental Material

**Supplementary Figure 1.**
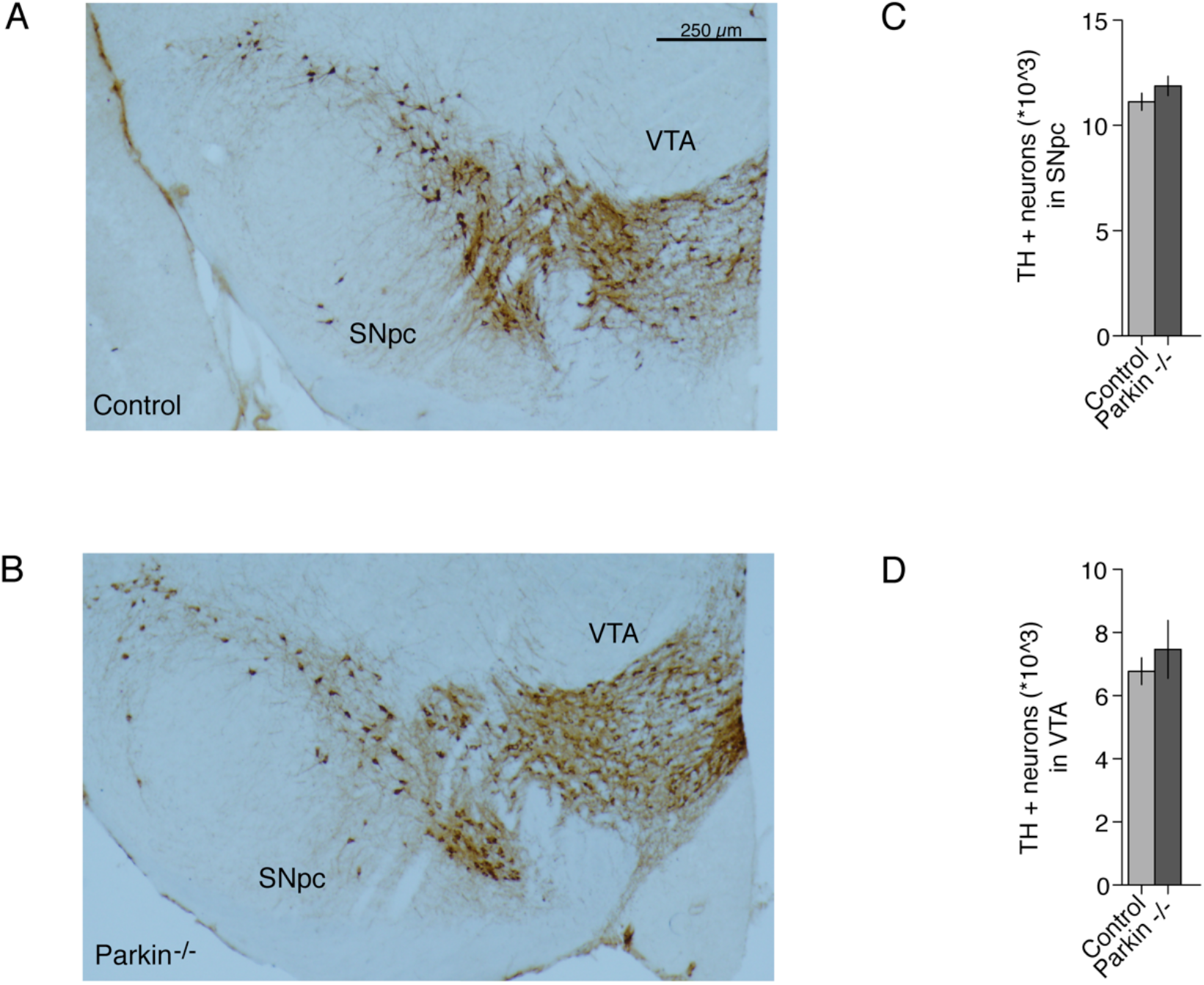
Stereological counting of Thyrosine Hydroxylase (TH) + cells in the substantia nigra pars compacta (SNpc) and ventral tegmental area (VTA). Example of TH-staining in a control (A) and parkin^−/−^ mouse (B). Bar graphs show mean +/− std of TH+ cell counts in SNpc (C) and (VTA) (D). Note the similar cell counts between the two experimental groups (n=3 animals per group).

**Supplementary Figure 2.**
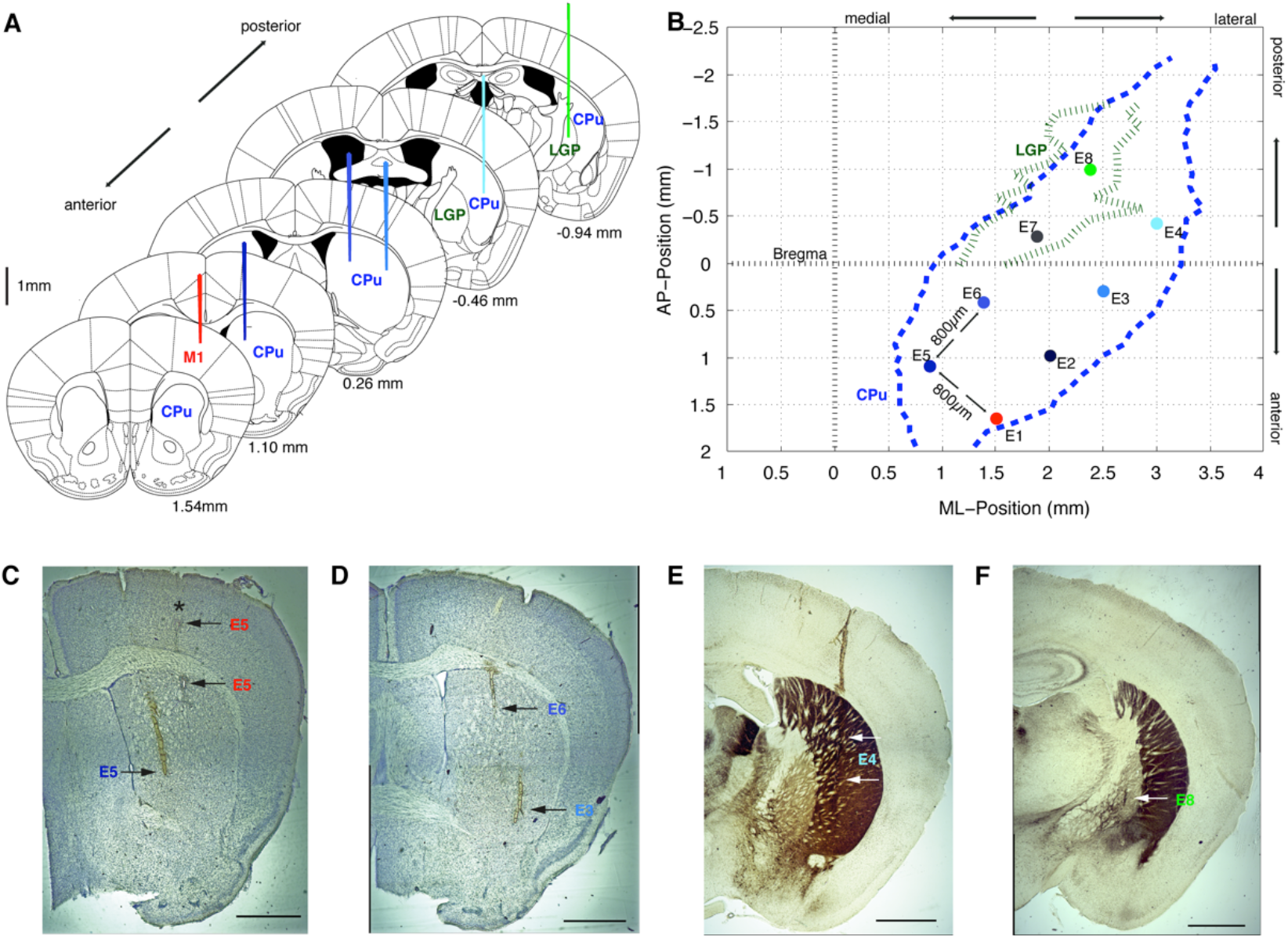
Recording setup and verification of recording sites. (A,B) Schematic illustration of recording setup. Example electrode positions for selected electrodes in the coronar plane are shown in (A) (Adapted from, The Mouse Brain in stereotactic coordinates’, Paxinos and Franklin, 2004). Note the position of one electrode in M1 of the motor cortex, several electrodes in CPu and one electrode in LGP (last section). Electrodes were organized in a grid of two electrode rows consisting of 4 electrodes. The distance between the rows was 800 µm as well as between the electrodes in one row (see grid in B). The grid was placed so that one electrode (E1) was positioned in the motor cortex M1, 1-2 electrodes in LGP (E7, E8) and all other electrodes E2-E6 in CPu. Approximate maximal borders of CPu and LGP were projected to surface and are shown in B. The blue dashed line represents borders of CPu and the green dashed line represents borders of LGP with respect to Bregma. Note that due to the grid configuration, if electrode 7 and 8 were located in LGP (identified by tonic high frequency discharge characteristics), the remaining electrodes were very likely positioned in CPu. In this recording setup, the depth of each electrode could be varied separately with a microdrive. Depth of 0 was defined as contact of each electrode with the cortical surface positioned under microscopic view. Moreover, slightly shifted positions were applied during the course of the experiment to get different recording positions (AP/ML) in one experiment. (D-F) Example coronar histology sections (50 µm) with corresponding electrode trajectories shown in A and B. Sections were alternating stained with Nissl (D,C) or Acetylcholinesterase (ACH) (E,F) using standard protocols. The scale bars indicate 1000 µm. Pictures were taken with a microscope (Carl Zeiss Microimaging, 2.5x magnification) using AxioVision software and Fiji Software for post hoc merging of pictures. (C) Example section approximately 1.1 mm anterior from Bregma. Arrows near the cortex show lesions of the cortical electrode E1 positioned approximately 1.5 mm anterior from Bregma. Lesions were performed at 800 and 1800 µm depth with respect to surface. The star indicates the recording position of the cortical electrode at 500 µm depth. The more medial arrow points at an electrode trace corresponding to E5 located in CPu. (D) Example section approximately 0.26 mm anterior from Bregma showing two striatal trajectories. The more medial trajectory probably corresponds to E6, while the more lateral trajectory corresponds to E3. (E) Example section approximately −0.46 posterior from Bregma showing LGP and CPu. The section shows a striatal trajectory corresponding to E4. (F) Example section approximately −0.94 mm posterior from Bregma showing the placement of the pallidal electrode E8 in LGP.

**Supplementary Figure 3.**
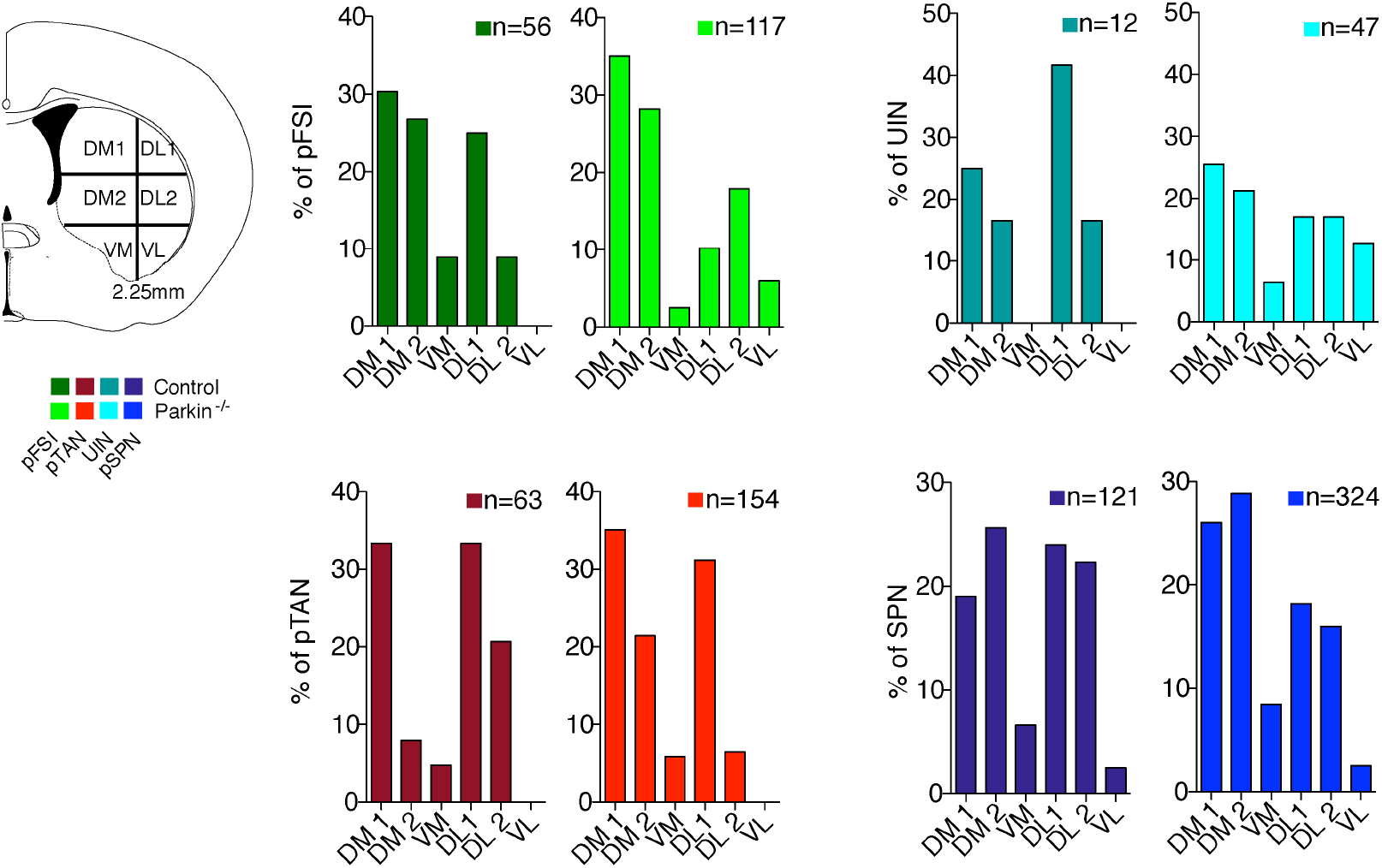
Distribution of recording sites for all striatal neuronal subpopulations. Left panel, Coronar brain slice (adapted from the mouse brain atlas of Paxinos and Franklin, 2004) with spatial subdivisions of the striatum used in this study. The striatum was divided into 3 subregions on each side along an idealized striatal midline. Dorsomedial 1 (DM1), dorsomedial 2 (DM2), ventromedial (VM) regions were located medial to the striatal midline (< 2.25 mm lateral to the whole brain midline). In contrast, dorsolateral 1 (DL1), dorsolateral 2 (DL2) and ventrolateral (VL) striatal territories were > 2.25mm lateral to whole brain midline. Furthermore, striatal subterritories were further subdivided according to depth below cortical surface: DM1/DL1: < 3 mm, DM2/DL2 between 3-4 mm, VM/VL > 4 mm. Right panels, Frequency distribution of striatal neuronal subtypes according to sampling site. Note the similar sampling/prevalence of different striatal neuron types in both experimental groups. Color code as within the main manuscript. Bar graphs with darker colors represent data from controls and brighter colors refer to data from parkin mutant mice.

**Supplementary Figure 4.**
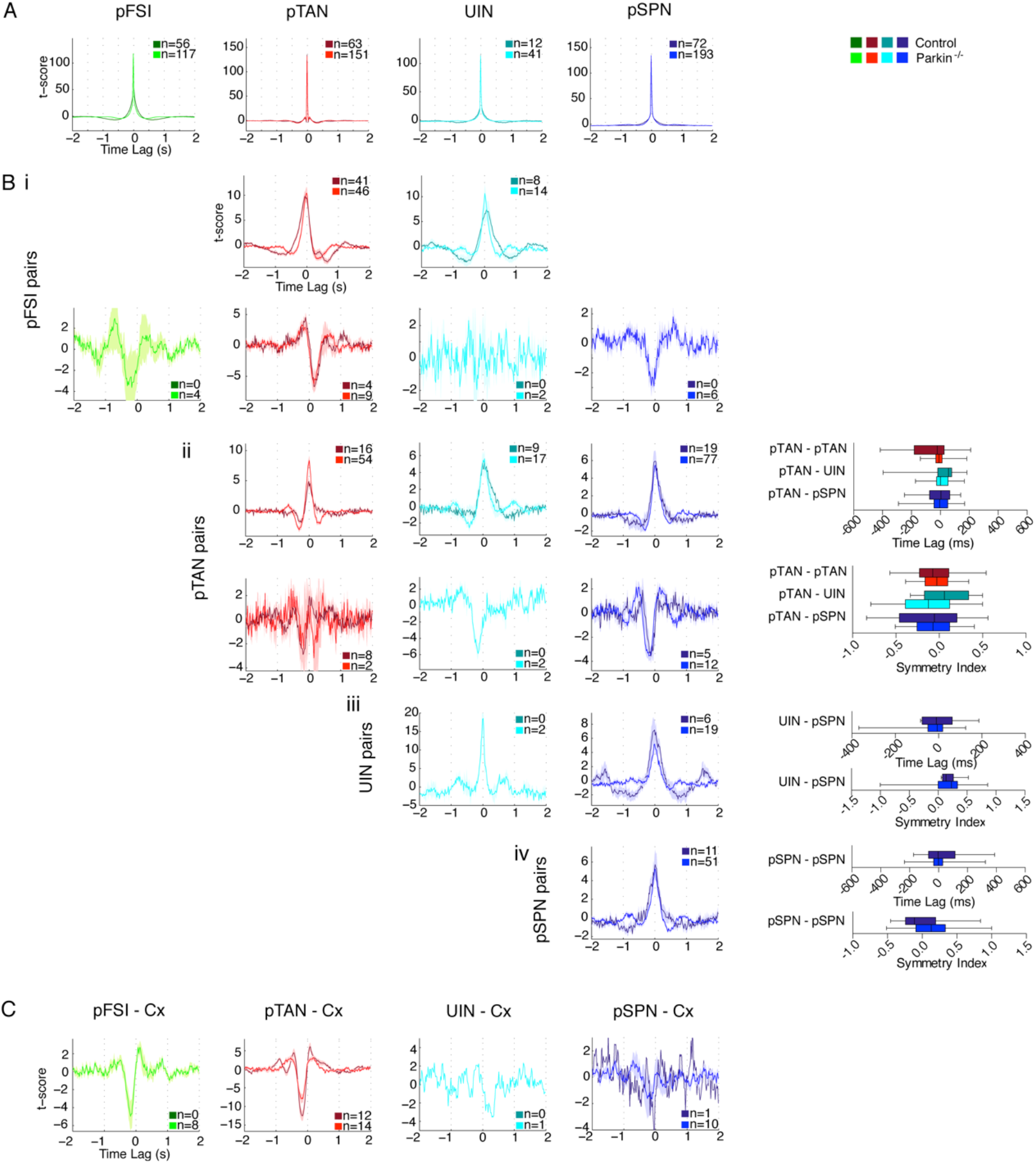
Comprehensive overview on auto- and cross-correlation histograms for all subgroups of striatal neurons. (A) Auto-correlation histograms. (Bi-iv) Left, Matrix of mean cross-correlation histograms derived from simultaneously recorded pairs of different striatal neuron types for both experimental groups. Color code is the same as in Figure 3 of the main text. Shading is SEM. Boxplots on the right provide an overview on statistical descriptors (time lag and symmetry index of the center peak) derived from the depicted cross-correlations. For further information, the reader is referred to the result section and Figure 3. The pairs shown here show no differences between the control and the parkin^−/−^ group. Whiskers of box plots show 5-95 percentile. (Bi) Cross-correlation histograms of positively and negatively correlated pFSIs with other striatal neuron types not shown in Figure 3. (Bii) Positively (top row) and negatively correlated (bottom row) pTAN pairs with other striatal neuron classes. (Biii) Positively correlated UIN pairs with other striatal neuron classes (no negatively correlated pairs available). (Biv) pSPN-pSPN pairs (no negative correlated pairs available). (C) T-scored cross-correlation histograms of significantly negative correlated cortico-striatal pairs (positive correlated pairs are provided in Figure 4 of the main text).

**Supplementary Figure 5.**
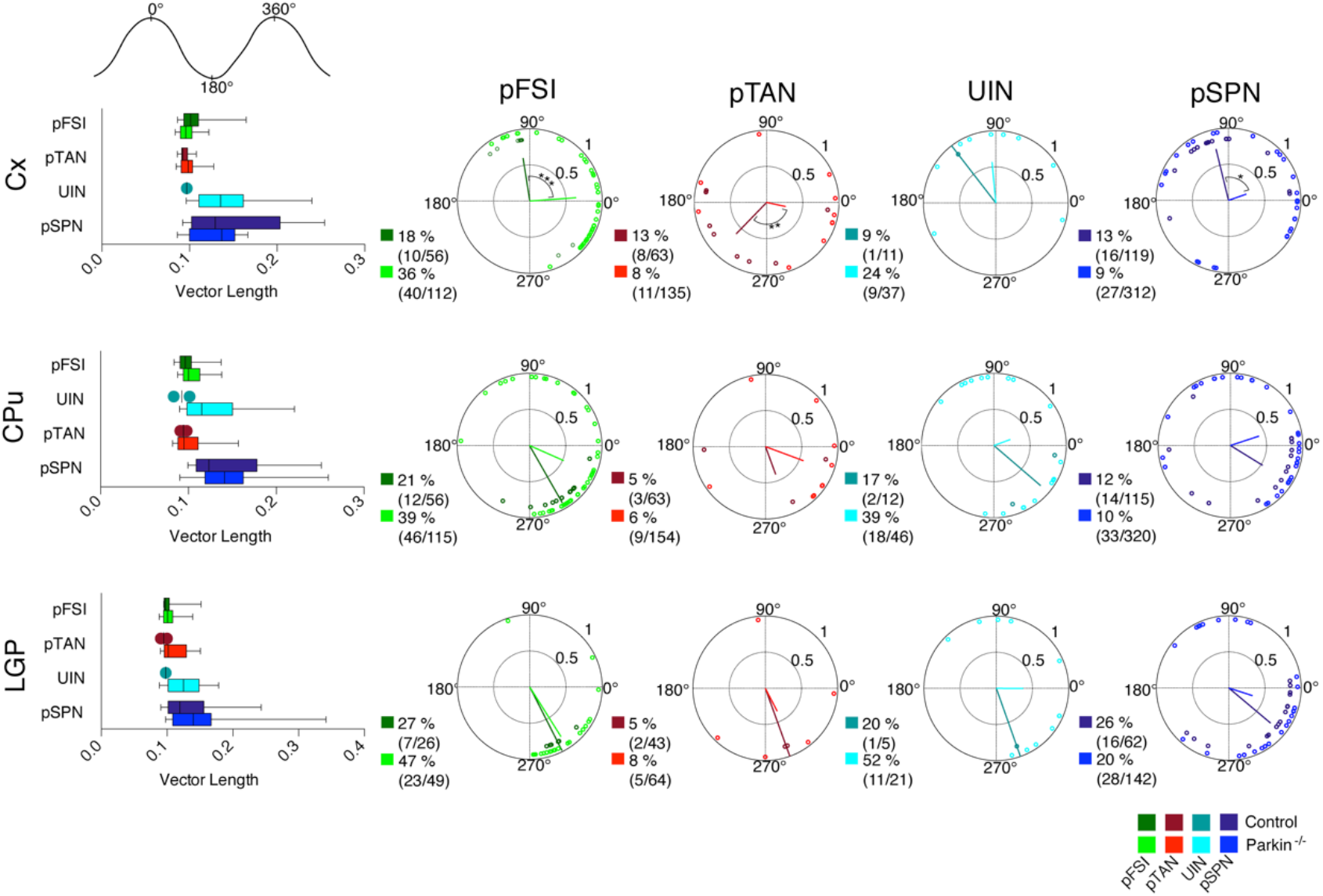
Phase locking of striatal neurons to ongoing LFP oscillations in the beta frequency range (20-30Hz). In both experimental groups, all striatal neuron subclasses of neurons and LGP single units were strongly and significantly phase locked to the cortical (top row), striatal (middle row) and pallidal (bottom row) LFPs filtered in the beta frequency range (i.e. between 20-30 Hz). Left panels, pairwise comparisons of the vector length (an rate-normalized indicator of phase locking strength) for all neuron types and neurons that were significantly phase-locked to beta oscillatory LFP activities, respectively. The circular plots in right panels show mean phase angles for neurons that were significantly locked to LFPs from motor cortex (top row), CPu (middle row) and LGP (bottom row), respectively. A dot represents the mean phase angle of an individual neuron. Vector length is derived from the variance of all mean angles. A phase angle of 180° corresponds to the negative peak of the LFP oscillation. For each group, both percentage and absolute number of significantly locked neurons are stated. Note that even though the prevalence of significant beta locking is generally much lower compared to spike field locking in the delta frequency range, the percentage of significantly locked pFSIs and UIN is higher in the parkin^−/−^ group in comparison to controls (see Figure 9 in the main text). As above, darker and brighter colors in all panels represent data from controls and parkin^−/−^ mice, respectively. Whiskers of box plots denote the 5-95 percentile. (*) p<0.05 before correction, *p<0.05, ** p<0.01, *** p<0.001

**Supplementary Table 1.** Descriptive statistics of striatal auto-correlations. Values for the maximal t-score of the peak of autocorrelation (+/− 2 s, bin size 20 ms) for all available putative striatal neurons with spike trains with more than 300 spikes. Differences were tested using MWUt and p-values are shown after FDR-correction.

| Neuron type | Experimental group | Number of neurons (>300 spikes) | Maximal T-score |  | Peak Duration (ms) |  |
| --- | --- | --- | --- | --- | --- | --- |
| | | | Mean ± Std | p-value | Mean ± Std | p-value |
| pFSI | control | 56 | 69.15 ± 11.78 | 0.75 | 500.9 ± 176.9 | 0.0004 |
| | parkin -/- | 117 | 69.65 ± 15.44 | | 393.2 ± 196.9 | |
| pTAN | control | 63 | 86.59 ± 9.324 | 0.75 | 234.8 ± 180.8 | 0.20 |
| | parkin -/- | 151 | 85.99 ± 13.05 | | 244.6 ± 136.1 | |
| UIN | control | 12 | 65.92 ± 29.88 | 0.88 | 522.9 ± 238.5 | 0.70 |
| | parkin -/- | 42 | 62.44 ± 29.88 | | 555.1 ± 333.8 | |
| pSPN | control | 72 | 77.64 ± 15.02 | 0.75 | 666.6 ± 278.3 | 4.4 E-06 |
| | parkin -/- | 194 | 79.89 ± 18.17 | | 500 ± 229.7 | |
| LGP SUAs | control | 55 | 56.61 ± 23.34 | 0.68 | 352.5 ± 329.1 | 0.20 |
| | parkin -/- | 55 | 63.19 ± 26.15 | | 483.1 ± 390.8 | |

**Supplementary Table 2.** Descriptive statistics of striatal, cortico-striatal, cortico-pallidal and striato-pallidal cross-correlations. The table shows the total number of significantly correlated cross-correlations (defined as 3 neighbored bins with a t-score >2) for significantly positive and negative correlated pairs. The absolute maximal t- score of cross-correlations expressing a significant peak is compared between the two experimental groups using MWUt. Note the significant higher absolute t-score of pFSI-pFSI in the control group indicating decreased pFSI-coupling in parkin^−/−^ mice. Moreover, the pair distance of simultaneous recorded striatal neuron pairs was compared between the two experimental groups, revealing no significant difference using MWUt.

| Cross-correlation Pair | Experimental Group | % significantly correlated | % positively correlated | % negatively correlated | Maximal t-score |  | Pair Distance (mm) |  |
| --- | --- | --- | --- | --- | --- | --- | --- | --- |
|  |  |  |  |  | (only significant) | p-value | (all available pairs) | p-value |
| pFSI – pFSI | control | 96.8 % (30/31) | 96.8 % (30/31) |  | 29.5 ± 14.8 | 9.1E-05 | 1.6 ± 0.6 | 0.59 |
|  | parkin -/- | 85.4 % (35/41) | 75.6 % (31/41) | 9.8 % (4/41) | 15.2 ± 8.5 |  | 1.5 ± 0.6 |  |
| pFSI – pTAN | control | 95.7 % (45/47) | 76.6 % (41/47) | 19.1 % (4/47) | 15.6 ± 7.2 | 0.88 | 1.5 ± 0.6 | 0.27 |
|  | parkin -/- | 73.3 % (55/75) | 61.3 % (46/75) | 12 % (9/75) | 16.0 ± 9.3 |  | 1.4 ± 0.6 |  |
| pFSI – UIN | control | 66.7 % (8/12) | 66.7 % (8/12) |  | 9.1 ± 5.8 | 0.77 | 1.4 ± 0.5 | 0.99 |
|  | parkin -/- | 61.5 % (16/26) | 53.8 % (14/26) | 7.7 % (2/26) | 12.6 ± 7.4 |  | 1.3 ± 0.4 |  |
| pFSI – pSPN | control | 76.6 % (36/47) | 76.6 % (36/47) |  | 11.4 ± 6.4 | 0.77 | 1.6 ± 0.6 | 0.53 |
|  | parkin -/- | 64.5 % (69/107) | 58.9 % (63/107) | 5.6 % (6/107) | 10.0 ± 5.2 |  | 1.5 ± 0.6 |  |
| pTAN – pTAN | control | 96 % (24/25) | 64 % (16/25) | 32% (8/25) | 12.0 ± 4.0 | 0.08 | 1.5 ± 0.6 | 0.18 |
|  | parkin -/- | 83.6 % (56/67) | 80.6 % (54/67) | 3 % (2/67) | 17.7 ± 9.5 |  | 1.2 ± 0.4 |  |
| pTAN – UIN | control | 75 % (9/12) | 75 % (9/12) |  | 10.3 ± 5.1 | 0.88 | 1.4 ± 0.4 | 0.89 |
|  | parkin -/- | 67.9 % (19/28) | 60.7 % (17/28) | 7.1 % (2/28) | 10.6 ± 7.1 |  | 1.4 ± 0.5 |  |
| pTAN – pSPN | control | 55.8 % (24/43) | 44.2 % (19/43) | 11.6 % (5/43) | 11.1 ± 7.1 | 0.88 | 1.4 ± 0.5 | 0.62 |
|  | parkin -/- | 68.5 % (89/130) | 59.2 % (77/130) | 9.2 % (12/130) | 9.6 ± 4.3 |  | 1.4 ± 0.5 |  |
| UIN – UIN | control | 0 % (0/2) |  |  |  |  |  |  |
|  | parkin -/- | 40 % (2/5) | 40% (2/5) |  | 23.6 ± 7.4 |  | 1.4 ± 0.7 |  |
| UIN – pSPN | control | 60 % (6/10) | 60% (6/10) |  | 10.1 ± 4.9 | 0.77 | 1.6 ± 0.8 | 0.94 |
|  | parkin -/- | 65.5 % (19/29) | 65.5 % (19/29) |  | 8.3 ± 4.3 |  | 1.5 ± 0.7 |  |

**Supplementary Table 2. Descriptive statistics of striatal, cortico-striatal, cortico-pallidal and striato-pallidal cross-correlations.**
| Cross-correlation Pair | Experimental | % significantly | % positively | % negatively | Maximal t-score |  | Pair Distance (mm) |  |
| --- | --- | --- | --- | --- | --- | --- | --- | --- |
|  | Group | correlated | correlated | correlated | (only significant) | p-value | (all available pairs) | p-value |
| pSPN – pSPN | control | 52.2 % (12/23) | 47.8 % (11/23) | 4.3 % (1/23) | 9.1 ± 7.0 | 0.88 | 1.3 ± 0.5 | 0.53 |
|  | parkin -/- | 51.5 % (51/99) | 51.5 % (51/99) |  | 8.1 ± 4.1 |  | 1.5 ± 0.6 |  |
| pFSI – Cx | control | 96.1 % (49/51) | 96.1 % (49/51) |  | 19.7 ± 10.1 | 0.67 |  |  |
|  | parkin -/- | 91.9 % (102/111) | 84.7 % (94/111) | 7.2 % (8/111) | 19.4 ± 16.7 |  |  |  |
| pTAN – Cx | control | 96.7 % (59/61) | 77 % (47/61) | 19.7 % (12/61) | 11.6 ± 17.2 | 0.08 |  |  |
|  | parkin -/- | 92.4 % (121/131) | 81.7 % (107/131) | 10.7 % (14/131) | 18.4 ± 15.5 |  |  |  |
| UIN – Cx | control | 81.8 % (9/11) | 81.8 % (9/11) |  | 12.9 ± 8.3 | 0.91 |  |  |
|  | parkin -/- | 87.9 % (29/33) | 84.8 % (28/33) | 3% (1/33) | 15.3 ± 11.5 |  |  |  |
| pSPN – Cx | control | 87.5 % (56/61) | 85.9 % (55/61) | 1.6 % (1/61) | 12.2 ± 7.7 | 0.91 |  |  |
|  | parkin -/- | 87.3 % (165/189) | 82 % (155/189) | 5.3 % (10/189) | 12.3 ± 9.2 |  |  |  |
| Cx – LGP | control | 77.6 % (38/49) | 42.9 % (21/49) | 34.7 % (17/49) | 11.3 ± 9.0 | 0.90 |  |  |
|  | parkin -/- | 81.3 % (39/48) | 33.3 % (16/48) | 47.9 % (23/48) | 13.5 ± 10.5 |  |  |  |
| pFSI – LGP | control | 45 % (9/20) | 30 % (6/20) | 15 % (3/20) | 9.5 ± 3.7 | 1 |  |  |
|  | parkin -/- | 42.9 % (9/21) | 19 % (4/21) | 23.8 % (5/21) | 12.7 ± 9.8 |  |  |  |
| pTAN – LGP | control | 35.7 % (15/42) | 28.6 % (12/42) | 7.1 % (3/42) | 9.8 ± 3.9 | 0.60 |  |  |
|  | parkin -/- | 52.6 % (20/38) | 31.6 % (12/38) | 21.1 % (8/38) | 13.2 ± 8.0 |  |  |  |
| pSPN - LGP | control | 33.3 % (9/27) | 22.2 % (6/27) | 11.1 % (3/27) | 5.7 ± 2.7 | 0.20 |  |  |
|  | parkin -/- | 48.3 % (28/58) | 17.2 % (10/58) | 31 % (18/58) | 9.1 ± 5.7 |  |  |  |

**Supplementary Table 3.** Number of significant phase-locked MUA to cortical, striatal and pallidal LFP in the beta frequency range (20-30 Hz). The number of significantly phase-locked MUA in different target structures was compared between control and parkin^−/−^ mice using Fisher’s Exact test. The corresponding comparison of phase-locking strength (vector length) is shown in Figure 9 of the main manuscript. P-values were corrected for each LFP-structure investigated and FDR-corrected p-values are shown as well.

| LFP | MUA | Experimental group | Number of MUA significantly locked | Total number of MUA | % significantly locked MUA | p-value Fisher's Exact Test | FDR-corrected p-value |
| --- | --- | --- | --- | --- | --- | --- | --- |
| Cx-LFP | CPu-MUA | control | 196 | 360 | 54.4 | 0.0007 | 0.0014 |
|  |  | parkin -/- | 531 | 818 | 64.9 |  |  |
| Cx-LFP | LGP-MUA | control | 10 | 69 | 14.5 | 0.1150 | 0.1150 |
|  |  | parkin -/- | 22 | 88 | 25.0 |  |  |
| CPu-LFP | Cx-MUA | control | 55 | 79 | 69.6 | 0.0738 | 0.1108 |
|  |  | parkin -/- | 104 | 180 | 57.8 |  |  |
| CPu-LFP | CPu-MUA | control | 259 | 371 | 69.8 | 0.0064 | 0.0192 |
|  |  | parkin -/- | 671 | 868 | 77.3 |  |  |
| CPu-LFP | LGP-MUA | control | 19 | 61 | 31.1 | 0.6079 | 0.6079 |
|  |  | parkin -/- | 34 | 96 | 35.4 |  |  |
| LGP-LFP | Cx-MUA | control | 10 | 69 | 14.5 | 0.3615 | 0.3615 |
|  |  | parkin -/- | 22 | 88 | 25.0 |  |  |
| LGP-LFP | CPu-MUA | control | 151 | 212 | 71.2 | 0.0002 | 0.0007 |
|  |  | parkin -/- | 301 | 356 | 84.6 |  |  |
| LGP-LFP | LGP-MUA | control | 5 | 24 | 20.8 | 0.1021 | 0.1532 |
|  |  | parkin -/- | 9 | 19 | 47.4 |  |  |

## Literature

Adler A, Katabi S, Finkes I, Prut Y, Bergman H. 2013. Different correlation patterns of cholinergic and GABAergic interneurons with striatal projection neurons. Frontiers in systems neuroscience. 7:47.

Albin RL, Young AB, Penney JB. 1989. The functional anatomy of basal ganglia disorders. Trends in neurosciences. 12:366–375.

Arkinson C, Walden H. 2018. Parkin function in Parkinson’s disease. Science. 360:267–268.

Avila I, Parr-Brownlie LC, Brazhnik E, Castaneda E, Bergstrom DA, Walters JR. 2010. Beta frequency synchronization in basal ganglia output during rest and walk in a hemiparkinsonian rat. Experimental neurology. 221:307–319.

Bar-Gad I, Heimer G, Ritov Y, Bergman H. 2003. Functional correlations between neighboring neurons in the primate globus pallidus are weak or nonexistent. The Journal of neuroscience : the official journal of the Society for Neuroscience. 23:4012–4016.

Barroso-Flores J, Herrera-Valdez MA, Lopez-Huerta VG, Galarraga E, Bargas J. 2015. Diverse Short-Term Dynamics of Inhibitory Synapses Converging on Striatal Projection Neurons: Differential Changes in a Rodent Model of Parkinson’s Disease. Neural plasticity. 2015:573543.

Baumer T, Pramstaller PP, Siebner HR, Schippling S, Hagenah J, Peller M, Gerloff C, Klein C, Munchau A. 2007. Sensorimotor integration is abnormal in asymptomatic Parkin mutation carriers: a TMS study. Neurology. 69:1976–1981.

Beck MH, Haumesser JK, Kuhn J, Altschuler J, Kuhn AA, van Riesen C. 2016. Short- and long-term dopamine depletion causes enhanced beta oscillations in the cortico-basal ganglia loop of parkinsonian rats. Experimental neurology. 286:124–136.

Benjamini Y, Hochberg Y. 1995. Controlling the False Discovery Rate - a Practical and Powerful Approach to Multiple Testing. J Roy Stat Soc B Met. 57:289–300.

Bennett BD, Callaway JC, Wilson CJ. 2000. Intrinsic membrane properties underlying spontaneous tonic firing in neostriatal cholinergic interneurons. The Journal of neuroscience : the official journal of the Society for Neuroscience. 20:8493–8503.

Berens P. 2009. CircStat: A MATLAB Toolbox for Circular Statistics. J Stat Softw. 31:1–21.

Bergman H, Wichmann T, Karmon B, DeLong MR. 1994. The primate subthalamic nucleus. II. Neuronal activity in the MPTP model of parkinsonism. Journal of neurophysiology. 72:507–520.

Berke JD, Okatan M, Skurski J, Eichenbaum HB. 2004. Oscillatory entrainment of striatal neurons in freely moving rats. Neuron. 43:883–896.

Brazhnik E, Cruz AV, Avila I, Wahba MI, Novikov N, Ilieva NM, McCoy AJ, Gerber C, Walters JR. 2012. State-dependent spike and local field synchronization between motor cortex and substantia nigra in hemiparkinsonian rats. The Journal of neuroscience : the official journal of the Society for Neuroscience. 32:7869–7880.

Bronte-Stewart H, Barberini C, Koop MM, Hill BC, Henderson JM, Wingeier B. 2009. The STN beta-band profile in Parkinson’s disease is stationary and shows prolonged attenuation after deep brain stimulation. Experimental neurology. 215:20–28.

Brown P, Oliviero A, Mazzone P, Insola A, Tonali P, Di Lazzaro V. 2001. Dopamine dependency of oscillations between subthalamic nucleus and pallidum in Parkinson’s disease. The Journal of neuroscience : the official journal of the Society for Neuroscience. 21:1033–1038.

Buhmann C, Binkofski F, Klein C, Buchel C, van Eimeren T, Erdmann C, Hedrich K, Kasten M, Hagenah J, Deuschl G, Pramstaller PP, Siebner HR. 2005. Motor reorganization in asymptomatic carriers of a single mutant Parkin allele: a human model for presymptomatic parkinsonism. Brain : a journal of neurology. 128:2281–2290.

Burkhardt JM, Constantinidis C, Anstrom KK, Roberts DC, Woodward DJ. 2007. Synchronous oscillations and phase reorganization in the basal ganglia during akinesia induced by high-dose haloperidol. The European journal of neuroscience. 26:1912–1924.

Chen H, Lei H, Xu Q. 2018. Neuronal activity pattern defects in the striatum in awake mouse model of Parkinson’s disease. Behavioural brain research. 341:135–145.

Chu Y, Morfini GA, Langhamer LB, He Y, Brady ST, Kordower JH. 2012. Alterations in axonal transport motor proteins in sporadic and experimental Parkinson’s disease. Brain : a journal of neurology. 135:2058–2073.

Corti O, Lesage S, Brice A. 2011. What genetics tells us about the causes and mechanisms of Parkinson’s disease. Physiological reviews. 91:1161–1218.

Damodaran S, Evans RC, Blackwell KT. 2014. Synchronized firing of fast-spiking interneurons is critical to maintain balanced firing between direct and indirect pathway neurons of the striatum. Journal of neurophysiology. 111:836–848.

Dauer W, Przedborski S. 2003. Parkinson’s disease: mechanisms and models. Neuron. 39:889–909.

Dawson TM, Ko HS, Dawson VL. 2010. Genetic animal models of Parkinson’s disease. Neuron. 66:646–661.

DeBoer P, Abercrombie ED, Heeringa M, Westerink BH. 1993. Differential effect of systemic administration of bromocriptine and L-dopa on the release of acetylcholine from striatum of intact and 6-OHDA-treated rats. Brain research. 608:198–203.

Degos B, Deniau JM, Chavez M, Maurice N. 2009. Chronic but not acute dopaminergic transmission interruption promotes a progressive increase in cortical beta frequency synchronization: relationships to vigilance state and akinesia. Cerebral cortex. 19:1616–1630.

Edgerton JR, Jaeger D. 2011. Dendritic sodium channels promote active decorrelation and reduce phase locking to parkinsonian input oscillations in model globus pallidus neurons. The Journal of neuroscience : the official journal of the Society for Neuroscience. 31:10919–10936.

Elia AE, Del Sorbo F, Romito LM, Barzaghi C, Garavaglia B, Albanese A. 2014. Isolated limb dystonia as presenting feature of Parkin disease. Journal of neurology, neurosurgery, and psychiatry. 85:827–828.

Feingold J, Gibson DJ, DePasquale B, Graybiel AM. 2015. Bursts of beta oscillation differentiate postperformance activity in the striatum and motor cortex of monkeys performing movement tasks. Proceedings of the National Academy of Sciences of the United States of America. 112:13687–13692.

Fino E, Venance L. 2010. Spike-timing dependent plasticity in the striatum. Frontiers in synaptic neuroscience. 2:6.

Fries P, Reynolds JH, Rorie AE, Desimone R. 2001. Modulation of oscillatory neuronal synchronization by selective visual attention. Science. 291:1560–1563.

Gerfen CR, Baimbridge KG, Miller JJ. 1985. The neostriatal mosaic: compartmental distribution of calcium-binding protein and parvalbumin in the basal ganglia of the rat and monkey. Proceedings of the National Academy of Sciences of the United States of America. 82:8780–8784.

Gittis AH, Hang GB, LaDow ES, Shoenfeld LR, Atallah BV, Finkbeiner S, Kreitzer AC. 2011. Rapid target-specific remodeling of fast-spiking inhibitory circuits after loss of dopamine. Neuron. 71:858–868.

Gittis AH, Leventhal DK, Fensterheim BA, Pettibone JR, Berke JD, Kreitzer AC. 2011. Selective inhibition of striatal fast-spiking interneurons causes dyskinesias. The Journal of neuroscience : the official journal of the Society for Neuroscience. 31:15727–15731.

Gittis AH, Nelson AB, Thwin MT, Palop JJ, Kreitzer AC. 2010. Distinct roles of GABAergic interneurons in the regulation of striatal output pathways. The Journal of neuroscience : the official journal of the Society for Neuroscience. 30:2223–2234.

Goldberg MS, Fleming SM, Palacino JJ, Cepeda C, Lam HA, Bhatnagar A, Meloni EG, Wu N, Ackerson LC, Klapstein GJ, Gajendiran M, Roth BL, Chesselet MF, Maidment NT, Levine MS, Shen J. 2003. Parkin-deficient mice exhibit nigrostriatal deficits but not loss of dopaminergic neurons. The Journal of biological chemistry. 278:43628–43635.

Grunewald A, Kasten M, Ziegler A, Klein C. 2013. Next-generation phenotyping using the parkin example: time to catch up with genetics. JAMA neurology. 70:1186–1191.

Heimer G, Bar-Gad I, Goldberg JA, Bergman H. 2002. Dopamine replacement therapy reverses abnormal synchronization of pallidal neurons in the 1-methyl-4-phenyl-1,2,3,6-tetrahydropyridine primate model of parkinsonism. The Journal of neuroscience : the official journal of the Society for Neuroscience. 22:7850–7855.

Hernandez LF, Kubota Y, Hu D, Howe MW, Lemaire N, Graybiel AM. 2013. Selective effects of dopamine depletion and L-DOPA therapy on learning-related firing dynamics of striatal neurons. The Journal of neuroscience : the official journal of the Society for Neuroscience. 33:4782–4795.

Hilker R, Klein C, Ghaemi M, Kis B, Strotmann T, Ozelius LJ, Lenz O, Vieregge P, Herholz K, Heiss WD, Pramstaller PP. 2001. Positron emission tomographic analysis of the nigrostriatal dopaminergic system in familial parkinsonism associated with mutations in the parkin gene. Annals of neurology. 49:367–376.

Hjorth J, Blackwell KT, Kotaleski JH. 2009. Gap junctions between striatal fast-spiking interneurons regulate spiking activity and synchronization as a function of cortical activity. The Journal of neuroscience : the official journal of the Society for Neuroscience. 29:5276–5286.

Holt GR, Softky WR, Koch C, Douglas RJ. 1996. Comparison of discharge variability in vitro and in vivo in cat visual cortex neurons. Journal of neurophysiology. 75:1806–1814.

Ikarashi Y, Takahashi A, Ishimaru H, Arai T, Maruyama Y. 1997. Relations between the extracellular concentrations of choline and acetylcholine in rat striatum. Journal of neurochemistry. 69:1246–1251.

Itier JM, Ibanez P, Mena MA, Abbas N, Cohen-Salmon C, Bohme GA, Laville M, Pratt J, Corti O, Pradier L, Ret G, Joubert C, Periquet M, Araujo F, Negroni J, Casarejos MJ, Canals S, Solano R, Serrano A, Gallego E, Sanchez M, Denefle P, Benavides J, Tremp G, Rooney TA, Brice A, Garcia de Yebenes J. 2003. Parkin gene inactivation alters behaviour and dopamine neurotransmission in the mouse. Human molecular genetics. 12:2277–2291.

Karain B, Xu D, Bellone JA, Hartman RE, Shi WX. 2015. Rat globus pallidus neurons: functional classification and effects of dopamine depletion. Synapse. 69:41–51.

Kaufman L, Rousseeuw PJ. 1990. Finding groups in data an introduction to cluster analysis. New York etc.: Wiley.

Khan NL, Brooks DJ, Pavese N, Sweeney MG, Wood NW, Lees AJ, Piccini P. 2002. Progression of nigrostriatal dysfunction in a parkin kindred: an [18F]dopa PET and clinical study. Brain : a journal of neurology. 125:2248–2256.

Kita H, Kita T. 2011. Role of Striatum in the Pause and Burst Generation in the Globus Pallidus of 6-OHDA-Treated Rats. Frontiers in systems neuroscience. 5:42.

Kitada T, Asakawa S, Hattori N, Matsumine H, Yamamura Y, Minoshima S, Yokochi M, Mizuno Y, Shimizu N. 1998. Mutations in the parkin gene cause autosomal recessive juvenile parkinsonism. Nature. 392:605–608.

Kitada T, Pisani A, Karouani M, Haburcak M, Martella G, Tscherter A, Platania P, Wu B, Pothos EN, Shen J. 2009. Impaired dopamine release and synaptic plasticity in the striatum of parkin−/− mice. Journal of neurochemistry. 110:613–621.

Klein C, Westenberger A. 2012. Genetics of Parkinson’s disease. Cold Spring Harbor perspectives in medicine. 2:a008888.

Kondabolu K, Roberts EA, Bucklin M, McCarthy MM, Kopell N, Han X. 2016. Striatal cholinergic interneurons generate beta and gamma oscillations in the corticostriatal circuit and produce motor deficits. Proceedings of the National Academy of Sciences of the United States of America. 113:E3159–3168.

Koos T, Tepper JM. 2002. Dual cholinergic control of fast-spiking interneurons in the neostriatum. The Journal of neuroscience : the official journal of the Society for Neuroscience. 22:529–535.

Kordower JH, Olanow CW, Dodiya HB, Chu Y, Beach TG, Adler CH, Halliday GM, Bartus RT. 2013. Disease duration and the integrity of the nigrostriatal system in Parkinson’s disease. Brain : a journal of neurology. 136:2419–2431.

Kramer ER, Aron L, Ramakers GM, Seitz S, Zhuang X, Beyer K, Smidt MP, Klein R. 2007. Absence of Ret signaling in mice causes progressive and late degeneration of the nigrostriatal system. PLoS biology. 5:e39.

Kramer ER, Knott L, Su F, Dessaud E, Krull CE, Helmbacher F, Klein R. 2006. Cooperation between GDNF/Ret and ephrinA/EphA4 signals for motor-axon pathway selection in the limb. Neuron. 50:35–47.

Kravitz AV, Freeze BS, Parker PR, Kay K, Thwin MT, Deisseroth K, Kreitzer AC. 2010. Regulation of parkinsonian motor behaviours by optogenetic control of basal ganglia circuitry. Nature. 466:622–626.

Kuhn AA, Kempf F, Brucke C, Gaynor Doyle L, Martinez-Torres I, Pogosyan A, Trottenberg T, Kupsch A, Schneider GH, Hariz MI, Vandenberghe W, Nuttin B, Brown P. 2008. High-frequency stimulation of the subthalamic nucleus suppresses oscillatory beta activity in patients with Parkinson’s disease in parallel with improvement in motor performance. The Journal of neuroscience : the official journal of the Society for Neuroscience. 28:6165–6173.

Kuhn AA, Trottenberg T, Kivi A, Kupsch A, Schneider GH, Brown P. 2005. The relationship between local field potential and neuronal discharge in the subthalamic nucleus of patients with Parkinson’s disease. Experimental neurology. 194:212–220.

Lachaux JP, Rodriguez E, Martinerie J, Varela FJ. 1999. Measuring phase synchrony in brain signals. Human brain mapping. 8:194–208.

Lansink CS, Goltstein PM, Lankelma JV, Pennartz CM. 2010. Fast-spiking interneurons of the rat ventral striatum: temporal coordination of activity with principal cells and responsiveness to reward. The European journal of neuroscience. 32:494–508.

Le Van Quyen M, Foucher J, Lachaux J, Rodriguez E, Lutz A, Martinerie J, Varela FJ. 2001. Comparison of Hilbert transform and wavelet methods for the analysis of neuronal synchrony. Journal of neuroscience methods. 111:83–98.

Leblois A, Meissner W, Bioulac B, Gross CE, Hansel D, Boraud T. 2007. Late emergence of synchronized oscillatory activity in the pallidum during progressive Parkinsonism. The European journal of neuroscience. 26:1701–1713.

Legendy CR, Salcman M. 1985. Bursts and recurrences of bursts in the spike trains of spontaneously active striate cortex neurons. Journal of neurophysiology. 53:926–939.

Lemaire N, Hernandez LF, Hu D, Kubota Y, Howe MW, Graybiel AM. 2012. Effects of dopamine depletion on LFP oscillations in striatum are task- and learning-dependent and selectively reversed by L-DOPA. Proceedings of the National Academy of Sciences of the United States of America. 109:18126–18131.

Levy R, Ashby P, Hutchison WD, Lang AE, Lozano AM, Dostrovsky JO. 2002. Dependence of subthalamic nucleus oscillations on movement and dopamine in Parkinson’s disease. Brain : a journal of neurology. 125:1196–1209.

Levy R, Hutchison WD, Lozano AM, Dostrovsky JO. 2000. High-frequency synchronization of neuronal activity in the subthalamic nucleus of parkinsonian patients with limb tremor. The Journal of neuroscience : the official journal of the Society for Neuroscience. 20:7766–7775.

Levy R, Hutchison WD, Lozano AM, Dostrovsky JO. 2002. Synchronized neuronal discharge in the basal ganglia of parkinsonian patients is limited to oscillatory activity. The Journal of neuroscience : the official journal of the Society for Neuroscience. 22:2855–2861.

Lucking CB, Durr A, Bonifati V, Vaughan J, De Michele G, Gasser T, Harhangi BS, Meco G, Denefle P, Wood NW, Agid Y, Brice A, French Parkinson’s Disease Genetics Study G, European Consortium on Genetic Susceptibility in Parkinson’s D. 2000. Association between early-onset Parkinson’s disease and mutations in the parkin gene. The New England journal of medicine. 342:1560–1567.

Luk KC, Sadikot AF. 2001. GABA promotes survival but not proliferation of parvalbumin-immunoreactive interneurons in rodent neostriatum: an in vivo study with stereology. Neuroscience. 104:93–103.

Madeo G, Martella G, Schirinzi T, Ponterio G, Shen J, Bonsi P, Pisani A. 2012. Aberrant striatal synaptic plasticity in monogenic parkinsonisms. Neuroscience. 211:126–135.

Madeo G, Schirinzi T, Martella G, Latagliata EC, Puglisi F, Shen J, Valente EM, Federici M, Mercuri NB, Puglisi-Allegra S, Bonsi P, Pisani A. 2014. PINK1 heterozygous mutations induce subtle alterations in dopamine-dependent synaptic plasticity. Movement disorders : official journal of the Movement Disorder Society. 29:41–53.

Mallet N, Le Moine C, Charpier S, Gonon F. 2005. Feedforward inhibition of projection neurons by fast-spiking GABA interneurons in the rat striatum in vivo. The Journal of neuroscience : the official journal of the Society for Neuroscience. 25:3857–3869.

Mallet N, Micklem BR, Henny P, Brown MT, Williams C, Bolam JP, Nakamura KC, Magill PJ. 2012. Dichotomous organization of the external globus pallidus. Neuron. 74:1075–1086.

Mallet N, Pogosyan A, Marton LF, Bolam JP, Brown P, Magill PJ. 2008. Parkinsonian beta oscillations in the external globus pallidus and their relationship with subthalamic nucleus activity. The Journal of neuroscience : the official journal of the Society for Neuroscience. 28:14245–14258.

Mallet N, Pogosyan A, Sharott A, Csicsvari J, Bolam JP, Brown P, Magill PJ. 2008. Disrupted dopamine transmission and the emergence of exaggerated beta oscillations in subthalamic nucleus and cerebral cortex. The Journal of neuroscience : the official journal of the Society for Neuroscience. 28:4795–4806.

Martella G, Platania P, Vita D, Sciamanna G, Cuomo D, Tassone A, Tscherter A, Kitada T, Bonsi P, Shen J, Pisani A. 2009. Enhanced sensitivity to group II mGlu receptor activation at corticostriatal synapses in mice lacking the familial parkinsonism-linked genes PINK1 or Parkin. Experimental neurology. 215:388–396.

Martinez WL, Martinez AR. 2005. Exploratory data analysis with MATLAB. Boca Raton, FL: Chapman & Hall/CRC.

Martinez WL, Martinez AR, Solka JL. 2011. Exploratory data analysis with MATLAB. Boca Raton, FL: CRC Press.

McCarthy MM, Moore-Kochlacs C, Gu X, Boyden ES, Han X, Kopell N. 2011. Striatal origin of the pathologic beta oscillations in Parkinson’s disease. Proceedings of the National Academy of Sciences of the United States of America. 108:11620–11625.

McGarry LM, Packer AM, Fino E, Nikolenko V, Sippy T, Yuste R. 2010. Quantitative classification of somatostatin-positive neocortical interneurons identifies three interneuron subtypes. Frontiers in neural circuits. 4:12.

Meka DP, Muller-Rischart AK, Nidadavolu P, Mohammadi B, Motori E, Ponna SK, Aboutalebi H, Bassal M, Annamneedi A, Finckh B, Miesbauer M, Rotermund N, Lohr C, Tatzelt J, Winklhofer KF, Kramer ER. 2015. Parkin cooperates with GDNF/RET signaling to prevent dopaminergic neuron degeneration. The Journal of clinical investigation.

Mitra PP, Bokil H. 2008. Observed brain dynamics. Oxford: Oxford University Press.

Mojena R. 1977. Hierarchical Grouping Methods and Stopping Rules - Evaluation. Comput J. 20:359–363.

Moll CK, Buhmann C, Gulberti A, Fickel U, Poetter-Nerger M, Westphal M, Gerloff C, Hamel W, Engel AK. 2015. Synchronized cortico-subthalamic beta oscillations in Parkin-associated Parkinson’s disease. Clinical neurophysiology : official journal of the International Federation of Clinical Neurophysiology.

Moshel S, Shamir RR, Raz A, de Noriega FR, Eitan R, Bergman H, Israel Z. 2013. Subthalamic nucleus long-range synchronization-an independent hallmark of human Parkinson’s disease. Frontiers in systems neuroscience. 7:79.

Nini A, Feingold A, Slovin H, Bergman H. 1995. Neurons in the globus pallidus do not show correlated activity in the normal monkey, but phase-locked oscillations appear in the MPTP model of parkinsonism. Journal of neurophysiology. 74:1800–1805.

Nolte G, Bai O, Wheaton L, Mari Z, Vorbach S, Hallett M. 2004. Identifying true brain interaction from EEG data using the imaginary part of coherency. Clinical neurophysiology : official journal of the International Federation of Clinical Neurophysiology. 115:2292–2307.

O’Malley KL. 2010. The role of axonopathy in Parkinson’s disease. Experimental neurobiology. 19:115–119.

Oostenveld R, Fries P, Maris E, Schoffelen JM. 2011. FieldTrip: Open source software for advanced analysis of MEG, EEG, and invasive electrophysiological data. Computational intelligence and neuroscience. 2011:156869.

Oyama G, Yoshimi K, Natori S, Chikaoka Y, Ren YR, Funayama M, Shimo Y, Takahashi R, Nakazato T, Kitazawa S, Hattori N. 2010. Impaired in vivo dopamine release in parkin knockout mice. Brain research. 1352:214–222.

Panicker N, Dawson VL, Dawson TM. 2017. Activation mechanisms of the E3 ubiquitin ligase parkin. The Biochemical journal. 474:3075–3086.

Parthasarathy HB, Graybiel AM. 1997. Cortically driven immediate-early gene expression reflects modular influence of sensorimotor cortex on identified striatal neurons in the squirrel monkey. The Journal of neuroscience : the official journal of the Society for Neuroscience. 17:2477–2491.

Pavese N, Khan NL, Scherfler C, Cohen L, Brooks DJ, Wood NW, Bhatia KP, Quinn NP, Lees AJ, Piccini P. 2009. Nigrostriatal dysfunction in homozygous and heterozygous parkin gene carriers: an 18F-dopa PET progression study. Movement disorders : official journal of the Movement Disorder Society. 24:2260–2266.

Paxinos G, Franklin KBJ. 2004. ≪The≫ mouse brain in stereotaxic coordinates. Amsterdam: Elsevier Academic.

Pisani A, Bernardi G, Ding J, Surmeier DJ. 2007. Re-emergence of striatal cholinergic interneurons in movement disorders. Trends in neurosciences. 30:545–553.

Planert H, Szydlowski SN, Hjorth JJ, Grillner S, Silberberg G. 2010. Dynamics of synaptic transmission between fast-spiking interneurons and striatal projection neurons of the direct and indirect pathways. The Journal of neuroscience : the official journal of the Society for Neuroscience. 30:3499–3507.

Prosperetti C, Di Giovanni G, Stefani A, Moller JC, Galati S. 2013. Acute nigro-striatal blockade alters cortico-striatal encoding: an in vivo electrophysiological study. Experimental neurology. 247:730–736.

Quiroga RQ, Nadasdy Z, Ben-Shaul Y. 2004. Unsupervised spike detection and sorting with wavelets and superparamagnetic clustering. Neural computation. 16:1661–1687.

Quiroga-Varela A, Walters JR, Brazhnik E, Marin C, Obeso JA. 2013. What basal ganglia changes underlie the parkinsonian state? The significance of neuronal oscillatory activity. Neurobiology of disease. 58:242–248.

Raff MC, Whitmore AV, Finn JT. 2002. Axonal self-destruction and neurodegeneration. Science. 296:868–871.

Ramanathan S, Hanley JJ, Deniau JM, Bolam JP. 2002. Synaptic convergence of motor and somatosensory cortical afferents onto GABAergic interneurons in the rat striatum. The Journal of neuroscience : the official journal of the Society for Neuroscience. 22:8158–8169.

Raz A, Frechter-Mazar V, Feingold A, Abeles M, Vaadia E, Bergman H. 2001. Activity of pallidal and striatal tonically active neurons is correlated in mptp-treated monkeys but not in normal monkeys. The Journal of neuroscience : the official journal of the Society for Neuroscience. 21:RC128.

Rivlin-Etzion M, Ritov Y, Heimer G, Bergman H, Bar-Gad I. 2006. Local shuffling of spike trains boosts the accuracy of spike train spectral analysis. Journal of neurophysiology. 95:3245–3256.

Salin P, Lopez IP, Kachidian P, Barroso-Chinea P, Rico AJ, Gomez-Bautista V, Coulon P, Kerkerian-Le Goff L, Lanciego JL. 2009. Changes to interneuron-driven striatal microcircuits in a rat model of Parkinson’s disease. Neurobiology of disease. 34:545–552.

Sassone J, Serratto G, Valtorta F, Silani V, Passafaro M, Ciammola A. 2017. The synaptic function of parkin. Brain : a journal of neurology. 140:2265–2272.

Savola JM, Virtanen R. 1991. Central alpha 2-adrenoceptors are highly stereoselective for dexmedetomidine, the dextro enantiomer of medetomidine. European journal of pharmacology. 195:193–199.

Scarffe LA, Stevens DA, Dawson VL, Dawson TM. 2014. Parkin and PINK1: much more than mitophagy. Trends in neurosciences. 37:315–324.

Scherfler C, Khan NL, Pavese N, Eunson L, Graham E, Lees AJ, Quinn NP, Wood NW, Brooks DJ, Piccini PP. 2004. Striatal and cortical pre- and postsynaptic dopaminergic dysfunction in sporadic parkin-linked parkinsonism. Brain : a journal of neurology. 127:1332–1342.

Schindelin J, Arganda-Carreras I, Frise E, Kaynig V, Longair M, Pietzsch T, Preibisch S, Rueden C, Saalfeld S, Schmid B, Tinevez JY, White DJ, Hartenstein V, Eliceiri K, Tomancak P, Cardona A. 2012. Fiji: an open-source platform for biological-image analysis. Nature methods. 9:676–682.

Schmitzer-Torbert NC, Redish AD. 2008. Task-dependent encoding of space and events by striatal neurons is dependent on neural subtype. Neuroscience. 153:349–360.

Schneider SA, Talelli P, Cheeran BJ, Khan NL, Wood NW, Rothwell JC, Bhatia KP. 2008. Motor cortical physiology in patients and asymptomatic carriers of parkin gene mutations. Movement disorders : official journal of the Movement Disorder Society. 23:1812–1819.

Sharott A, Doig NM, Mallet N, Magill PJ. 2012. Relationships between the firing of identified striatal interneurons and spontaneous and driven cortical activities in vivo. The Journal of neuroscience : the official journal of the Society for Neuroscience. 32:13221–13236.

Sharott A, Gulberti A, Hamel W, Koppen JA, Munchau A, Buhmann C, Potter-Nerger M, Westphal M, Gerloff C, Moll CKE, Engel AK. 2018. Spatio-temporal dynamics of cortical drive to human subthalamic nucleus neurons in Parkinson’s disease. Neurobiology of disease. 112:49–62.

Sharott A, Gulberti A, Zittel S, Tudor Jones AA, Fickel U, Munchau A, Koppen JA, Gerloff C, Westphal M, Buhmann C, Hamel W, Engel AK, Moll CK. 2014. Activity parameters of subthalamic nucleus neurons selectively predict motor symptom severity in Parkinson’s disease. The Journal of neuroscience : the official journal of the Society for Neuroscience. 34:6273–6285.

Sharott A, Magill PJ, Harnack D, Kupsch A, Meissner W, Brown P. 2005. Dopamine depletion increases the power and coherence of beta-oscillations in the cerebral cortex and subthalamic nucleus of the awake rat. The European journal of neuroscience. 21:1413–1422.

Sharott A, Moll CK, Engler G, Denker M, Grun S, Engel AK. 2009. Different subtypes of striatal neurons are selectively modulated by cortical oscillations. The Journal of neuroscience : the official journal of the Society for Neuroscience. 29:4571–4585.

Sharott A, Vinciati F, Nakamura KC, Magill PJ. 2017. A Population of Indirect Pathway Striatal Projection Neurons Is Selectively Entrained to Parkinsonian Beta Oscillations. The Journal of neuroscience : the official journal of the Society for Neuroscience. 37:9977–9998.

Tachibana Y, Iwamuro H, Kita H, Takada M, Nambu A. 2011. Subthalamo-pallidal interactions underlying parkinsonian neuronal oscillations in the primate basal ganglia. The European journal of neuroscience. 34:1470–1484.

Tankus A, Yeshurun Y, Fried I. 2009. An automatic measure for classifying clusters of suspected spikes into single cells versus multiunits. Journal of neural engineering. 6:056001.

Tepper JM, Bolam JP. 2004. Functional diversity and specificity of neostriatal interneurons. Current opinion in neurobiology. 14:685–692.

Tseng KY, Kasanetz F, Kargieman L, Riquelme LA, Murer MG. 2001. Cortical slow oscillatory activity is reflected in the membrane potential and spike trains of striatal neurons in rats with chronic nigrostriatal lesions. The Journal of neuroscience : the official journal of the Society for Neuroscience. 21:6430–6439.

Wichmann T, DeLong MR. 1996. Functional and pathophysiological models of the basal ganglia. Current opinion in neurobiology. 6:751–758.

Williams D, Tijssen M, Van Bruggen G, Bosch A, Insola A, Di Lazzaro V, Mazzone P, Oliviero A, Quartarone A, Speelman H, Brown P. 2002. Dopamine-dependent changes in the functional connectivity between basal ganglia and cerebral cortex in humans. Brain : a journal of neurology. 125:1558–1569.

Wilson CJ, Goldberg JA. 2006. Origin of the slow afterhyperpolarization and slow rhythmic bursting in striatal cholinergic interneurons. Journal of neurophysiology. 95:196–204.

Wiltschko AB, Pettibone JR, Berke JD. 2010. Opposite effects of stimulant and antipsychotic drugs on striatal fast-spiking interneurons. Neuropsychopharmacology : official publication of the American College of Neuropsychopharmacology. 35:1261–1270.

Yael D, Zeef DH, Sand D, Moran A, Katz DB, Cohen D, Temel Y, Bar-Gad I. 2013. Haloperidol-induced changes in neuronal activity in the striatum of the freely moving rat. Frontiers in systems neuroscience. 7:110.

Yamin HG, Stern EA, Cohen D. 2013. Parallel processing of environmental recognition and locomotion in the mouse striatum. The Journal of neuroscience : the official journal of the Society for Neuroscience. 33:473–484.

Zeitler M, Fries P, Gielen S. 2006. Assessing neuronal coherence with single-unit, multi-unit, and local field potentials. Neural computation. 18:2256–2281.

Zold CL, Ballion B, Riquelme LA, Gonon F, Murer MG. 2007. Nigrostriatal lesion induces D2-modulated phase-locked activity in the basal ganglia of rats. The European journal of neuroscience. 25:2131–2144.

Zold CL, Larramendy C, Riquelme LA, Murer MG. 2007. Distinct changes in evoked and resting globus pallidus activity in early and late Parkinson’s disease experimental models. The European journal of neuroscience. 26:1267–1279.

Ztaou S, Maurice N, Camon J, Guiraudie-Capraz G, Kerkerian-Le Goff L, Beurrier C, Liberge M, Amalric M. 2016. Involvement of Striatal Cholinergic Interneurons and M1 and M4 Muscarinic Receptors in Motor Symptoms of Parkinson’s Disease. The Journal of neuroscience : the official journal of the Society for Neuroscience. 36:9161–9172.

Zuurbier CJ, Emons VM, Ince C. 2002. Hemodynamics of anesthetized ventilated mouse models: aspects of anesthetics, fluid support, and strain. American journal of physiology Heart and circulatory physiology. 282:H2099–2105.

